# Specimen-tailored “lived” climate reveals precipitation onset and amount best predict specimen phenology, but only weakly predict estimated reproduction across a clade

**DOI:** 10.1101/2025.02.03.636077

**Authors:** Megan Bontrager, Samantha J. Worthy, Laura Leventhal, Julin A. Maloof, Jennifer R. Gremer, Johanna Schmitt, Sharon Y. Strauss

## Abstract

Herbarium specimens are broadly distributed in space and time, enabling investigation of climate impacts on phenology and fitness. We reconstructed specimen “lived” climate from knowledge of germination cues and collection dates for 14 annual species in the *Streptanthus* (s.l.) clade (Brassicaceae) to ask: Which climate attributes, including the timing of precipitation onset, best explain specimen phenological stage and estimated reproduction? We also asked whether climate effects on phenology and reproduction were evolutionarily conserved.

Precipitation amount and onset date, more than temperature, best predicted specimen phenology, but only weakly predicted reproduction. Earlier rainfall onset was associated with more phenologically advanced specimens, a relationship that showed phylogenetic signal. Few climate predictors explained variation in specimen reproduction. The lack of association between specimen reproduction and climate may arise from phenological shifts that buffer impacts of climate, interactions with other species, or challenges in estimating total reproduction from specimens.

Our results highlight the value of specimen-tailored growing season conditions for reconstructing climate, incorporating evolutionary relationships in assessing responses to climate, and the complexities of estimating fitness from specimens. For the latter, we propose supplemental herbarium collections and community science protocols to increase the utility of these data for understanding climate impacts on populations.

## Introduction

As climate change alters the environmental conditions that species experience, an urgent and open question is how climate affects plant performance and ultimately population stability. A number of studies have addressed how changing rainfall and temperature may affect plant growth, phenology, and reproduction in greenhouse and field experiments, and the results are often complex. In natural settings, diverse antagonists and mutualists may mediate plant responses to climate, contributing to the complexity of outcomes (González-Megías & Menéndez, 2012; Barnett & Facey, 2016; Wade *et al*., 2017). Herbarium specimens collected from natural plant populations provide valuable historical data and allow for insights into how climate conditions affect plant performance (Davis *et al*., 2015; Willis *et al*., 2017; Meineke *et al*., 2018) by integrating over biotic and abiotic environments. Since specimens are collected over large spatial scales and across diverse habitats, as well as over many years with varying climate conditions, analyses of specimen characteristics can be complementary to experimental approaches for understanding impacts of climate on plant phenology and fitness.

Numerous studies have analyzed phenological data from herbarium specimens and have revealed signatures of changing climate on phenological responses, as well as differences among species and regions in phenological sensitivity to climate variation (Primack *et al*., 2004; Miller-Rushing *et al*., 2006; Calinger *et al*., 2013; Heberling & Isaac, 2018; Jones & Daehler, 2018; Iler *et al*., 2019; DeLeo *et al*., 2020; Love & Mazer, 2021; Pearson *et al*., 2021; Willems *et al*., 2022; Dangremond *et al*., 2022; Ramirez-Parada *et al*., 2024). In contrast, only a few studies have analyzed measures of fitness or reproductive success from herbarium data, and to date, these studies have focused on integrating individual performance with species distribution models to investigate the climatic factors contributing to species range limits (Bontrager & Angert, 2016; Yim *et al*., 2024). Climate effects on reproductive success may, however, be more closely tied to population viability than are phenological shifts (Iler *et al*., 2019). Additionally, phenology and reproductive success are not just affected by the magnitude but also the timing of weather events (Steyn *et al*., 1996; Levine *et al*., 2011; Olliff-Yang & Ackerly, 2021; Martínez-Berdeja *et al*., 2023; Worthy *et al*., 2025). In western North America, recent studies have shown that rainfall onset is occurring later in fall and precipitation in fall is decreasing (Swain *et al*., 2018; Luković *et al*., 2021). These trends, layered onto increasing temperature trends over the past century (Pathak *et al*., 2018), are effectively increasing drought stress, shifting seasonal timing, and potentially decoupling temperature and precipitation patterns (Flint *et al*., 2013; Swain *et al*., 2018; Luković *et al*., 2021).

The timing of rainfall may have especially large effects on the growing season in Mediterranean ecosystems where rainfall occurs in only a few months of the year, beginning in the fall and ending with the onset of months-long drought in spring. Winter annual plant species, whose germination is triggered by fall rainfall, may suffer fitness impacts from shortened growing seasons with later-arriving rainfall (Steyn *et al*., 1996; Levine *et al*., 2011; Olliff-Yang & Ackerly, 2021; Martínez-Berdeja *et al*., 2023; Worthy *et al*., 2025). In addition to affecting growing season length, germination timing determines a plant’s “lived climate”: the realized climate niche experienced during its lifetime (Bontrager et al. 2025), including the seasonal cues regulating flowering and fruiting phenology. In a screenhouse experiment (where plants were exposed to ambient temperatures but watering regimes that simulated different timing of rainfall onset), later rainfall onset caused some native annual species to germinate later and subsequently accelerate their flowering phenology. This acceleration partially made up for fitness losses due to shortened growing seasons, but could not fully compensate for the effects of shorter growing seasons on reproductive fitness (Worthy *et al*., 2025); see also (Steyn *et al*., 1996; Levine *et al*., 2011; Olliff-Yang & Ackerly, 2021; Martínez-Berdeja *et al*., 2023; Zettlemoyer *et al*., 2024). In the field, effects of delayed rainfall onset also depend on rainfall distribution and total amounts—an early rainfall year, which could benefit plant reproduction by lengthening the growing season, might be disastrous if drought ensues (Iler *et al*., 2019), and mid-winter drought is becoming more common in the west (Hatchett *et al*., 2023).

Here, we ask how precipitation, temperature, and the timing of rainfall onset affect the phenological stage and estimated reproduction of herbarium specimens. We also ask whether the same climatic variables affect both of these reproductive properties. We use new methods that focus on reconstructing a tailored “lived climate” for each specimen from the estimated germination date in the fall until collection in the following spring or summer. We examined specimens of 14 annual species in the native *Streptanthus* (*s.l.*) complex (Brassicaceae) that germinate in the fall and complete reproduction in early to mid-summer. We studied this clade because we have extensive knowledge of its ecology (see below) as well as a robust evolutionary hypothesis (Cacho *et al*., 2014) and thus can also ask to what extent phenological and reproductive responses to climate reflect evolutionary history.

In this system, the timing and extent of rainfall affect phenology and fitness, and these effects vary across the clade and in relation to local climate. Our previous work provides some *a priori* expectations for how these species might respond to changing precipitation levels or timing in the field and also shows that evolutionary history of the clade may influence these responses (Pearse *et al*., 2022; Worthy *et al*., 2025). Our most recent work in this system suggests that though these species occupy diverse latitudes and elevations (Fig. S1), growing season climate conditions, especially climate water deficit (CWD) and temperature, are more constrained than expected, suggesting phylogenetic niche conservatism (Bontrager *et al*., 2025). In a common garden study of 11 of these annual *Caulanthus* and *Streptanthus* species, Pearse et al. (2022) found that higher-aridity *Caulanthus* species flowered relatively earlier than species from wetter habitats and could maintain reproduction at field-relevant extremely low water levels, but were unable to extend their flowering periods when ample water was provided. In contrast, taxa from relatively wetter Mediterranean climates flowered later and extended their flowering periods substantially with ample water, but were unable to reproduce at the lowest water levels. These results suggest that precipitation timing could be important to these species. A separate experiment manipulating the seasonal timing of rainfall showed that fitness generally declined with later rainfall onset, but that the magnitude of this effect varied among species; more mesic species accelerated flowering more than arid species in response to late rainfall arrival, and such compensatory plasticity resulted in smaller fitness declines (Worthy *et al*., 2025). Our prior experiments on germination timing, in which we manipulated timing of rainfall onset under ambient outdoor conditions, revealed that these species germinate at highest rates in the fall with early rainfall onset (Worthy *et al*., 2024). Taken *in toto*, our previous results indicate that some of these species could be vulnerable to climate change, in particular, to changes in rainfall timing.

Here, we follow up from this previous work by using customized growing season windows for each herbarium specimen to ask: 1) How does “lived” climate—precipitation, temperature and climatic water deficit—affect the phenology and estimated reproduction of herbarium specimens? 2) Does later onset of seasonal rainfall alter reproductive phenology or reduce estimated specimen reproduction? 3) Across species, are phenological and estimated reproduction responses to climate correlated and 4) Are responses to climate evolutionarily conserved across the clade? In other words, Are close relatives more similar in their reproductive responses to climate than more distantly related species, a pattern that might indicate constraints on evolution.

Lastly, in the process of collecting data from specimens, we recognized that additions to current collection protocols could make herbarium specimens even more valuable for asking questions especially about reproduction and population persistence. We propose supplementing current herbarium collection protocols with additional types of specimen collection strategies, and linkages to community science digital images (see also Heberling and Isaac 2018 ; Iler et al 2019). Ultimately, herbarium specimens are, and will continue to be, very useful for understanding the impacts of environmental change on natural populations and their long-term viability.

## Materials and Methods

### Study system, species selection, and image acquisition

*Streptanthus* (s.l.) is a Brassicaceae clade that arose in the southwestern desert of western North America (Cacho *et al*., 2014) and continued to speciate as species moved west and northward as part of what Raven and Axelrod (1978) termed the Madro-Tertiary flora (Cacho *et al*., 2014, 2021). The Streptanthus I subclade comprises approximately 40 species in the *Streptanthus* and *Caulanthus* genera that occupy a diversity of habitats, spanning desert ecosystems to mid-montane habitats (Cacho *et al*., 2014). They are primarily annual species and all share an affinity for open, bare, rocky or sandy habitat (Cacho & Strauss, 2014). These species encompass a remarkable diversity of range sizes, latitudinal ranges, and elevational distributions (Fig. S1). Thus, the clade occupies a variety of climatic conditions (Pearse *et al*., 2020; Cacho *et al*., 2021; Worthy *et al*., 2024), and within some of the more wide-ranging taxa, populations can span a large amount of intraspecific climatic variation (Gremer *et al*., 2020; Love & Mazer, 2021).

From the species in *Streptanthus* I clade (Cacho *et al*., 2014), we chose 14 annual species to include in this study. The selection of species was based on three criteria: 1) sampling subclades within this group, 2) sampling species from across the latitudinal and elevational space, and thus the range of climate space, that each species occupies, and 3) sampling species with what we hoped would be an adequate sample size of individuals (>40 specimens). The last criterion resulted in the exclusion of the most narrow-ranged endemic species, which comprise about 20% of the group.

To assess phenological stage and estimated reproduction at the time of collection, we counted and categorized reproductive structures on high resolution digital images of herbarium sheets available via the Consortium of California Herbaria specimen data portal (cch2.org). We supplemented these images with herbarium sheets from the California Academy of Sciences herbarium that we photographed ourselves. These efforts resulted in inventories of 50-186 herbarium sheets per species (average = 138). When >200 sheets were available for a given species, we focused our counting efforts on approximately 200 sheets, subsampling by county to maximize geographic coverage. Among these subsets, species differed qualitatively in the temporal distribution of specimens across years 1898-2016, however all species were represented by specimens collected across a wide range of years (Fig. S2). All data were collected at the individual plant level. Often, multiple plants were mounted on a single sheet (average of 2.4 specimens/sheet, range = 1-12; non-independence was addressed with random effect structure, see statistical methods). Before counting an individual, we checked for the presence of roots, rosette leaves or rosette leaf scars and only counted individuals that appeared to be entire plants.

### Counting and measuring specimens

Because machine learning has been shown to undercount reproductive structures of herbarium specimens (Love *et al*., 2021), a team of fifteen people counted and classified reproductive structures on 4602 individuals collected on 1939 specimen sheets. Each counter was trained on the same set of specimens by the same team member using an annotated guide to the different types of structures. Counts were spot-checked for consistency across species and counters. We used a custom plugin in ImageJ (modified from CellCounter) to count and categorize reproductive structures; this plugin made counting more accurate by allowing the counter to select structure types by name— e.g., buds, flowers, fruits—and place color coded counters on the images. These counts were then saved in .xml files for collation and spot-checking. Reproductive structures were classified in detail, including missing structures that we inferred to be buds, flowers, immature fruits and mature fruits, based on persistent empty pedicels and the position of those pedicels relative to other buds, flowers and fruits (see Table S1, Fig. S3, S4). “Empty” pedicels could arise from selective abortion of structures, natural damage before collection, or loss during specimen preparation. These inferred or unknown structures make up a relatively small part of our dataset (6.1% of 167,398 total structures counted). Further, because pedicels from lost or aborted fruits were once flowers whose pollen could sire seed on other plants (part of male fitness), we chose to include all pedicels in our reproductive structure count, even if they did not produce fruit. Some species have sterile buds forming a showy tuft at the top of the plant; the structures comprising this tuft were not counted. Species varied in the dates and phenological stages at which they were collected (Fig. S5).

### Georeferencing

Geographic data and locality descriptions for each herbarium sheet were downloaded from the Consortium of California Herbaria and checked for consistency with described collection sites. We used Google Earth, complemented by the USGS Historical Topo Map database (https://apps.nationalmap.gov/downloader/#/maps) and results from google searches of location descriptions. We added error radius distances when they were missing. These were estimated by the georeferencer as the distance at which the provided description would no longer be the most logical. When specimens were recent and the label indicated that coordinates were recorded in the field with a GPS, rather than by another party after specimen accession, we tended to assign a smaller error radius if missing (though in most cases these collections already had an error radius recorded). Specimens for which we could not determine coordinates with <=5 km error radius (n = 112) were not used (though the large majority of specimens had much more precise coordinates, mean = 1.1 km, median = 0.5 km).

### Estimating phenological stage and reproduction

To quantify phenological stage, we first calculated a weighted sum of reproductive structures on each plant (0*number of buds + 0.33*number of flowers + 0.66*number of immature fruits + 1*number of mature fruits). This weighted sum is then divided by the sum of all reproductive structures on a plant, and the resulting metric is our phenological stage variable. In this method, each reproductive structure was weighted by its position in a phenological progression from buds to fruits (with “early” structures getting lower weights and “later” structures getting higher weights). This is similar to the metric used in Love *et al*. (2019), but scaled so that a binomial response distribution can be used. To analyse these data we use binomial generalized linear models, in which the weighted sum of reproductive structures (rounded to the nearest whole number) is used as the number of “successes” and the total number of reproductive structures is used as the number of “trials” (for all other model details see Statistical analyses: Q1). Binomial models are suitable for proportional data and allow us to maintain information about the number of structures from which phenological stage was estimated. Based on our metric, a specimen with a phenological stage of 0.1 would mostly have buds, while a specimen with a stage of 0.9 would be mostly in mature fruit. To estimate reproduction, we used the sum of all reproductive structures, generously assuming that every reproductive structure would make a fruit, and bearing in mind that flowers contribute to male fitness of plants.

In our statistical models, we accounted for the length of time that a specimen was able to grow using the day of year that a specimen was collected, as well as estimated germination dates, (see below); we expected that individuals that grew for longer periods of time had more time to advance in phenology and to reproduce, regardless of the climate conditions they experienced. The collection day-of-year variable was calculated as the number of days starting from September 1 in the fall prior to collection until the collection date the following spring/summer. Collection day-of-year (DOY) was included in every model as it was frequently positively related to our response variables of phenology and total reproduction (Figs. S6, S7).

### Estimating growing season climate

In Mediterranean ecosystems, precipitation starts in the fall, typically peaks in the winter and is very rare in the summer months (Di Castri & Mooney, 2012). For annual species in Mediterranean climates, fall precipitation of sufficient quantity stimulates germination and species complete their life cycles in a single season before surviving the summer drought as seeds (Keeley, 1991). Prior experiments on germination behavior in the*Streptanthus* clade (Worthy *et al*., 2024) and other winter annual plants have found that most species have their greatest germination during early fall rain events (Levine *et al*., 2008; Hänel & Tielbörger, 2015; Martínez-Berdeja *et al*., 2023), which can start as early as September. To estimate the timing of germination, we identified relevant rain events, which we considered to be when a total of 25 mm or more precipitation fell in a single event over one or more days. This threshold precipitation amount was chosen based on prior work showing that 25 mm of rain was sufficient to trigger germination in *Streptanthus tortuosus* (Gremer *et al*., 2020), and is the same threshold used in other systems, including desert annuals (Went, 1949; Tevis Jr., 1958; Beatley, 1974; Schwinning & Sala, 2004). We used this threshold in two ways, which we describe in detail below. With monthly climate data, we used the first month preceding collection with >25mm rain as the start of the specimen growing season and calculated other climate conditions using this month as the starting month. We also used data available at a daily resolution to look specifically at the effects of this rainfall timing on phenology and estimated reproduction; in this case the date of a rain event is going into the model directly rather than just delineating the start of the window over which other climate variables are averaged or summed.

Monthly climate data were extracted from the California Basin Characterization Model (Flint & Flint, 2014). We used monthly data for each specimen beginning with the first month after August with >25 mm precipitation (totalled over the month) and ending with the average collection month across all specimens of a species. We used the average collection month of each species, rather than the specific collection month of each specimen, to avoid confounding our climate characterizations with collection timing. We extracted maximum temperature (tmx), minimum temperature (tmn), precipitation (ppt), and climatic water deficit (CWD). We calculated average temperature (monthly (min + max)/2), summed CWD, and summed precipitation for the time period preceding collection of the specimen.

We also used climate data available at a daily resolution from the PRISM database (PRISM Climate Group, Oregon State University, https://prism.oregonstate.edu, accessed 14 Dec 2023). We used these daily data to estimate germination timing at a finer temporal scale than the monthly data allowed, using the same >25mm threshold. We omitted eight specimens from this subset of the data because there was no rain event >25mm until after March 1, which is no longer a plausible germination time (Worthy *et al*. 2024). Unfortunately, daily climate records are only available starting in 1981, and thus only specimens collected after 1981 could be used for analyses that included daily-resolution putative germination timing as a predictor (n = 828).

### Statistical analyses

#### Q1: How does “lived” climate affect the phenology and estimated reproduction of herbarium specimens?

The climate variables we used were average temperature, summed precipitation, and summed CWD, averaged or summed across the estimated growing season to capture what each specimen experienced in its lifetime. CWD is an overall measure of drought stress, and incorporates temperature, precipitation, soil water holding capacity, and aspect. Because CWD is derived in part from temperature and precipitation, it is correlated with both of these variables (positively with temperature and negatively with precipitation). We fit two different models for each response variable (phenological stage and total reproduction): one that included specimen-experienced temperature and precipitation together, and another with CWD alone. The model with CWD alone allows us to explore the effects of drought stress specifically, while the model with temperature and precipitation allows us to see the relative importance of these two climate variables for plant reproduction.

To estimate the effects of climate on phenology, we fit generalized linear mixed effects models with a binomial response distribution (see “Estimating phenological stage and reproduction” above) and a logit link function for each species separately. To estimate the effects of climate on estimated reproduction, we fit generalized linear mixed effects models with a negative binomial (gamma-Poisson) response distribution and a log link function, again using data from each species separately. In both cases, models were fit with the Bayesian regression package brms (Bürkner, 2017) and included a covariate of collection day-of-year. We could imagine scenarios in which a quadratic relationship might better explain the relationship between total reproduction and climate; for example, under low precipitation, total reproduction could be reduced by drought, while at high precipitation, total reproduction could be reduced by inter- or intraspecific competition (Hänel & Tielbörger, 2015). To explore these possibilities, we checked whether a quadratic function better explained the relationship between total reproduction and climate variables than a linear one; in the vast majority of cases, quadratic terms for temperature, precipitation, and CWD were not significant (the exceptions were *C. cooperi*, which had highest reproduction at intermediate CWD, *C. anceps*, which had highest reproduction at intermediate precipitation and high/low temperatures, *S. insignis*, which had highest reproduction at intermediate temperatures) and thus only linear relationships were tested in the models reported here.

For analyses related to questions 1 and 2, we included a group-level (random) effect of collection location, which was generated by placing buffers around each collection locality with a radius of 2.5 km (or their coordinate uncertainty, if larger than 2.5 km) and assigning points with overlapping buffers to the same group level (this method allowed “daisy chain” effects where two points without overlapping buffers were joined by a common neighbor). Multiple herbarium sheets are sometimes collected at the same location (e.g., to deposit in different herbaria) and so the non-independence of the specimens on these sheets is also represented in this effect.

Predictors (collection day-of-year and climate variables) were centered and scaled before analysis. We used a weak normal prior on betas, the default Student-t prior on the intercept, the default bounded Student-t prior on the standard deviation of group-level intercepts, and the default inverse-gamma prior on the shape parameter. All parameters were assessed for convergence (Rhat < 1.01). Parameter estimates and convergence metrics for all regression models are provided in Appendix 1 (see online supporting information). We interpret effects to be different from zero when the 95% credible interval of the estimate does not include 0. We visualised predicted regression slopes and credible intervals using the ggemmeans() function from the package ggeffects (Lüdecke, 2018). For each model for each species, we checked whether predictor variables were correlated with each other. In all cases except three, correlation coefficients were <0.6, in the remaining three they were <0.7.

#### Q2: Does later onset of seasonal rainfall alter phenology or reduce estimated reproduction?

*A priori,* we predicted from results of our previous screenhouse experiments that late rainfall would result in later reproduction (represented by lower phenology scores, controlling for collection date), and ultimately would decrease fitness (Worthy *et al*., 2025). Conversely, early rainfall should lead to earlier reproduction (higher phenology score) and increase fitness by lengthening the growing season, independent of rainfall amount and temperature. To test these hypotheses, we used the subset of specimens for which we had estimated germination date at the daily scale (see Methods: Estimating growing season climate). Using this subset of the data, we ran models with average temperature, total precipitation, and collection day-of-year as described for Q1 but added estimated germination date as an additional predictor. All other model details are as described for Q1. We again checked each species for correlations between the four predictors used. Most pairs of predictors had correlation coefficients below 0.6, six species had one to three pairs of predictors with correlation coefficients >0.6 and <0.76.

#### Q3: Across species, are phenological and estimated reproduction responses to climate variables correlated?

Plasticity in phenology in response to climate may be adaptive and may buffer fitness from the effects of climate variation. Species with phenologies that respond to climate variation may be buffered from effects of climate on total reproduction. In contrast, species that are unable or do not buffer climate variation via phenological timing—that is, have more fixed phenologies regardless of the climate they experience—should be more affected by climate variation (Worthy *et al*., 2025). Across species, we tested for correlations between slopes of the relationships between species’ phenological and reproductive responses to each climate variable using phylogenetic independent contrasts. We implemented these with the pic() function in the R package ape (Paradis & Schliep, 2019) and the phylogeny from Cacho et al. (2014).

#### Q4: Are responses to climate evolutionarily conserved across the clade?

To evaluate whether there might be evolutionary constraints in how species respond to climate (temperature, precipitation, CWD, and the timing of the first rainfall event) in their phenology and estimated reproduction, we tested for phylogenetic signal in effect size and direction from the models above. Phylogenetic signal indicates that close relatives are more similar in the direction and degree to which they respond to climate than more distantly related species. For this, we used Blomberg’s K (Blomberg *et al*., 2003), calculated with the phylosig function in the phytools package (Revell, 2012). We used the estimated error in our model effect sizes as the sampling error. We estimated the significance of phylogenetic signal, i.e. whether the amount of signal exceeds the quantity expected by random chance, using 10,000 randomizations where effect sizes were randomized across the tips of the phylogeny and K was repeatedly calculated. A p-value for this null hypothesis of no phylogenetic signal was then calculated as the fraction of randomizations with equal or higher K values than our observed estimate of K (Revell & Harmon, 2022). A non-significant K value (p > 0.05) suggests that phenology and estimated reproduction responses to climate are randomly distributed across the phylogeny. When significant (p < 0.05), we then tested whether the estimate of Blomberg’s K was significantly greater than one by comparing our observed estimate of K to a null distribution of 10,000 K values generated under Brownian motion, a null model for evolution (Revell & Harmon, 2022). P-values were calculated by counting the proportion of times the simulated values of K were greater than our observed estimate of K. K has an expected value of one under evolution by Brownian motion (Revell & Harmon, 2022), whereas values of K significantly greater than one indicate that species tend to have more similar phenology and estimated reproduction responses to climate than expected based on a Brownian motion model of evolution, i.e. more evolutionary constraint. Significant effects of phylogenetic relationships on the size and direction of climate effects might also indicate climate adaptation during species divergence.

## Results

### Q1: How does “lived” climate affect the phenology and estimated reproduction of herbarium specimens?

#### Phenology

We found effects of total precipitation, average temperature, and climatic water deficit (CWD) experienced by specimens on the phenological stage of the specimen (Figs. 1, 2, S8, S9). The direction and magnitude of effects varied among species and climate variables. Precipitation amounts significantly affected phenology for 10 species: increased annual precipitation significantly delayed phenology in eight species, slightly accelerated phenology in two species, and had no effect on four species (Fig. 1a, b), all while taking into account effects of average temperature and collection timing. Eight species adjusted phenological stage in response to temperature, with seven accelerating phenology with warmer temperatures, and one species delaying phenology with warmer temperatures (Figs. 1b, S8a). Seven species also significantly responded to CWD, albeit more idiosyncratically than to temperature or precipitation, with four species slightly accelerating phenology, and three slightly delaying phenology with increasing CWD (Figs. 1c, S8b). These analyses all included collection day-of-year as a covariate to account for the timing of the end of the specimen’s life, which together with germination date determines the total amount of time the specimen had to reach its phenological stage (Figs. S6a, S7a). Precipitation overall had stronger effects than temperature or CWD on phenology, based on the frequency of significant slopes and the magnitude of effect sizes relating phenology to climate variables.

**Figure 1.**
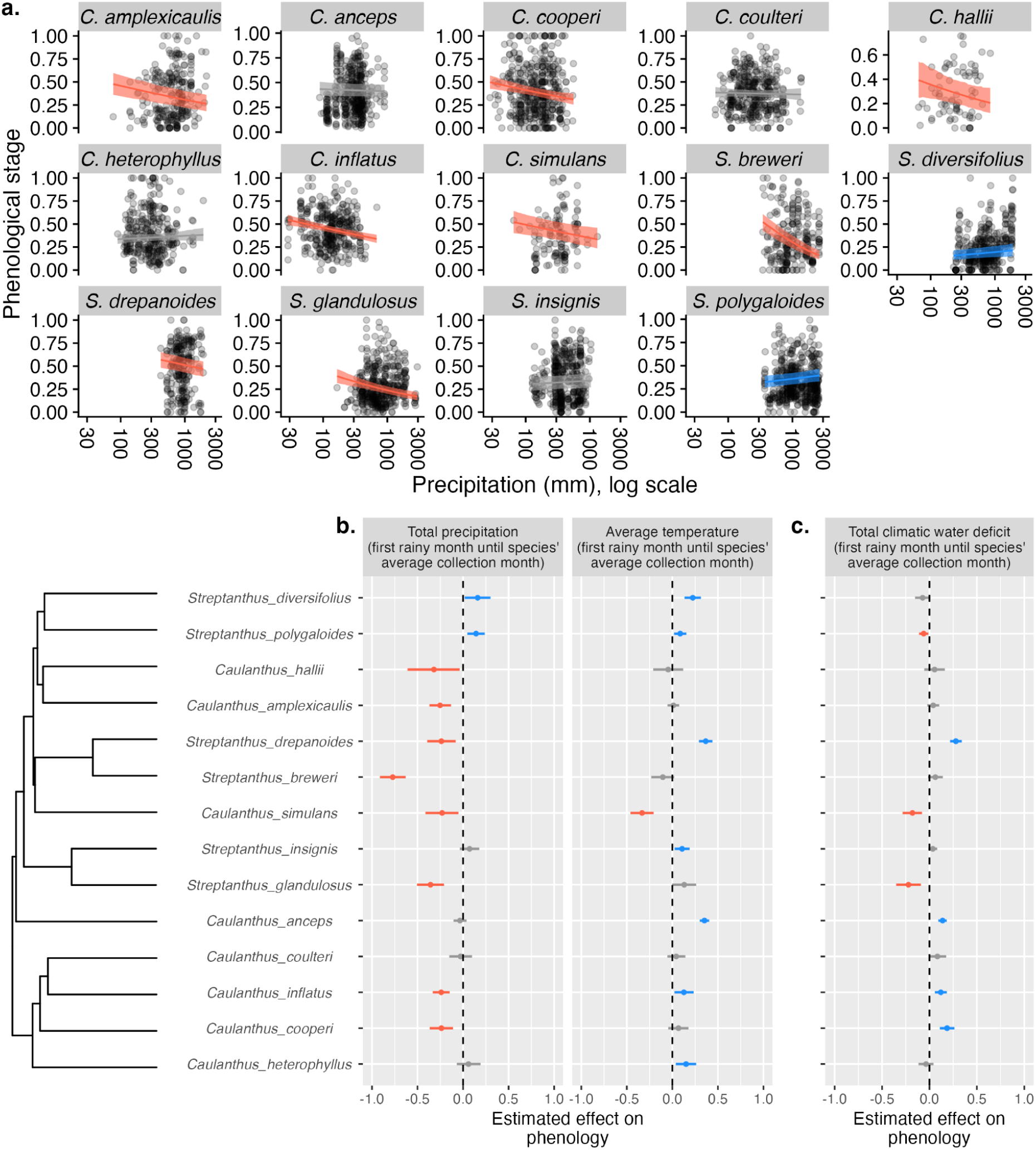
(a) Effects of precipitation on specimen phenological stage for 14 focal species. Regression slopes from models that include temperature and collection day-of-year (see Fig S6). Eight species had significantly more advanced phenology with less precipitation (red lines), while two species had delayed phenology with less precipitation (blue lines); gray lines are not significant. Similar plots for temperature and CWD are in Fig S10. (b) Mean slope with 95% credible interval of the regression of precipitation and temperature (in the same model) and (c) climatic water deficit on the phenological stage of specimens. The precipitation column in (b) represents the slopes depicted in (a). None of these responses to climate showed significant phylogenetic signal (K<1.0, P>0.4 in all cases; Table S3).

**Figure 2.**
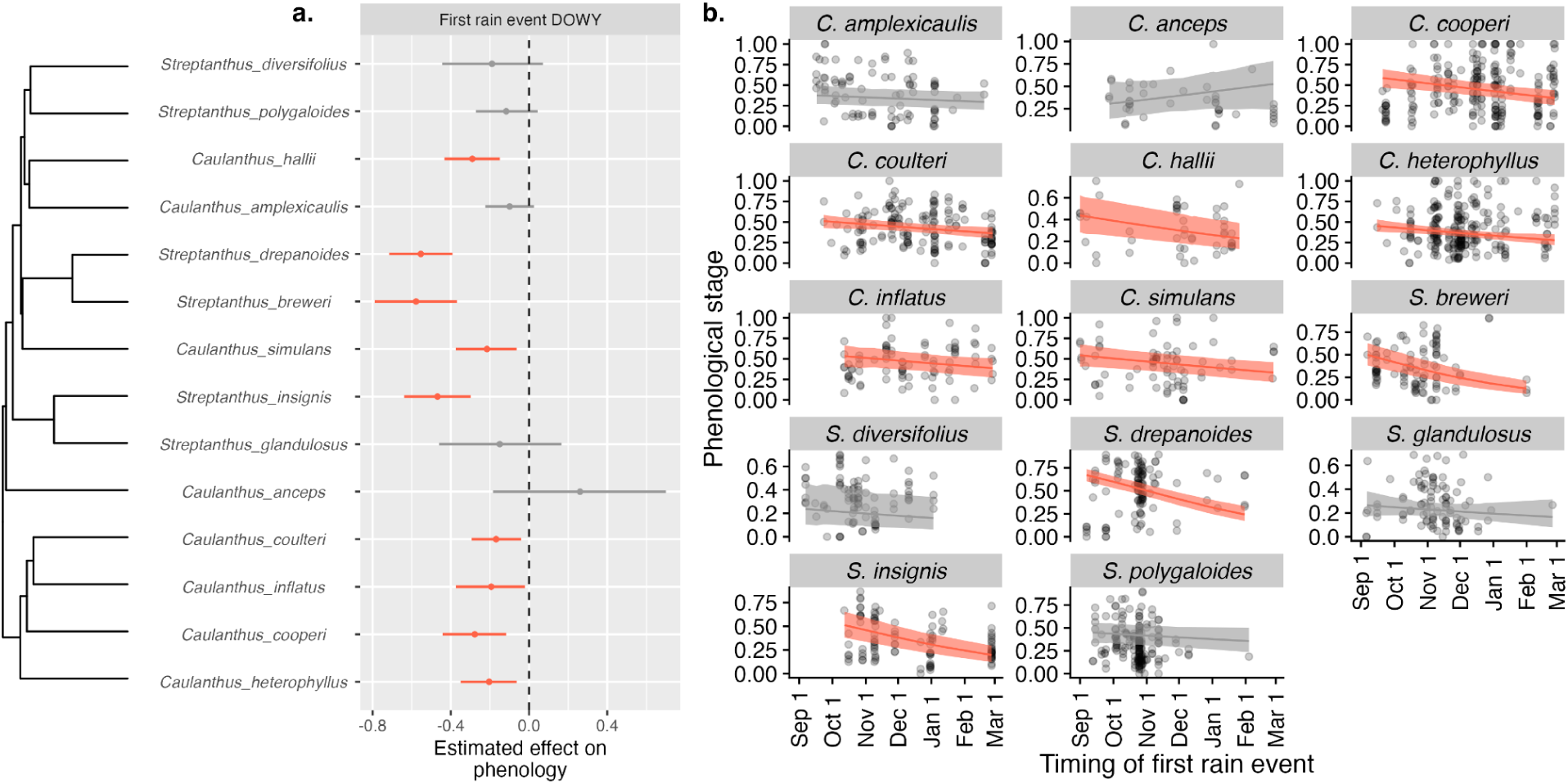
Effects of the timing of germination-triggering rain events on phenology of herbarium specimens. In (a) red point ranges indicate species for which later estimated germination timing delayed phenology. Grey point ranges show effects not different from zero. In (b) these slopes are plotted over the raw data points; phenological scores range from 0 (entirely in bud) to 1 (entirely mature fruit). Collection day-of-year, total precipitation and average temperature were also included in the model as predictors and these effects are shown in Fig. S9.

#### Estimated reproduction

Specimens collected later in the season had greater estimated reproduction for eight of fourteen species—the remaining six species showed no effect of collection day (Fig. S7b). Increased precipitation had a negative effect on the number of reproductive structures for five of the species—perhaps counterintuitively in California’s mediterranean climate and considering results from Pearse *et al*. (2020)—and a positive effect on two of the species (Fig. 3ab). As with phenology, temperature had fewer impacts on reproduction than did precipitation, with only two species showing significant responses (Figs. 3b, S10a). Higher temperatures were associated with lower reproduction for *S. diversifolius,* and were positively related to reproduction for *S. insignis*. Our separate model examining the relationships between CWD and estimated reproduction also showed varying effect sizes and directions (Figs. 3c, S10b). In four species increased CWD had positive effects on total reproduction and in two species increased CWD had negative effects on total reproduction.

**Figure 3.**
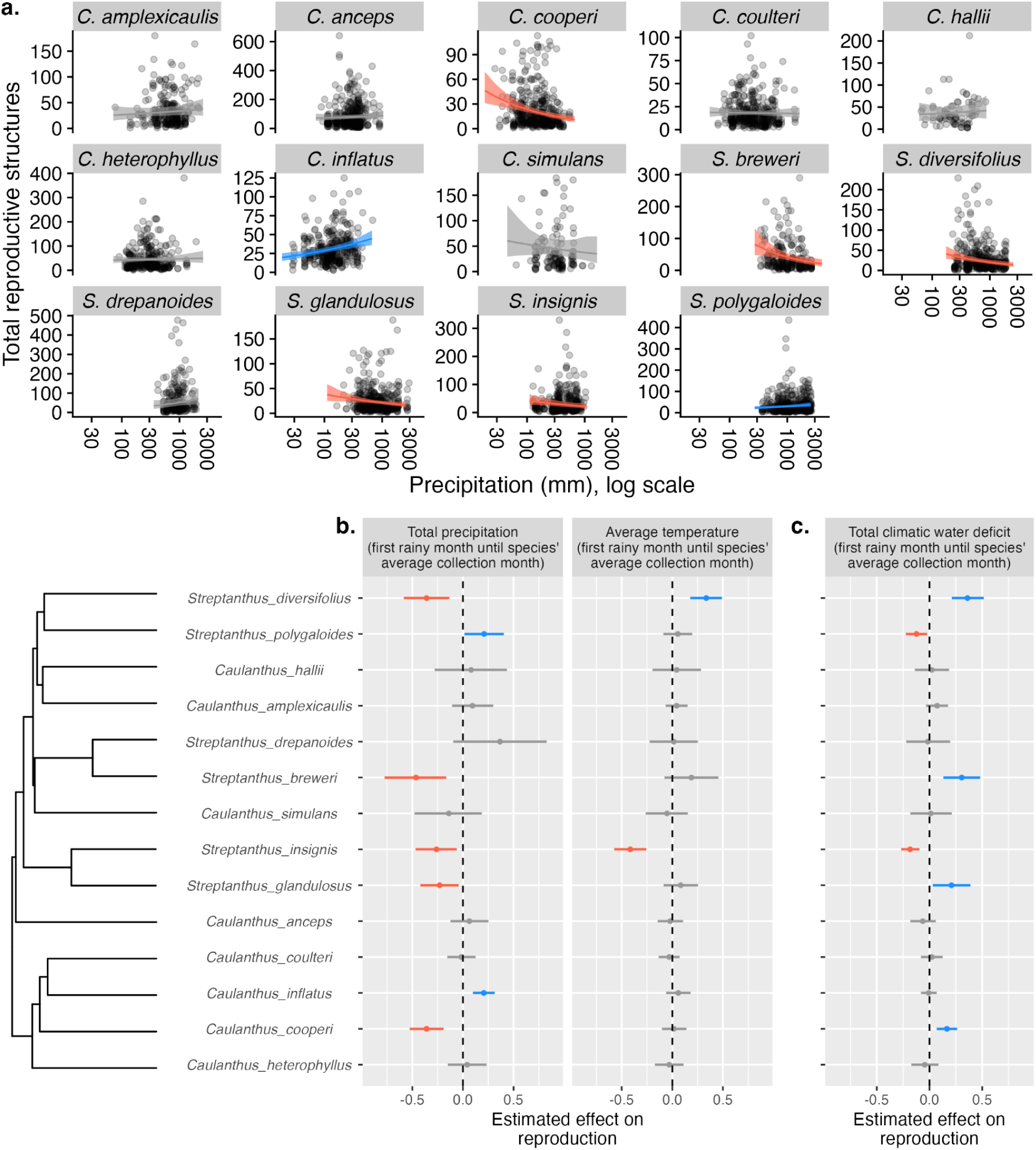
(a) Effects of precipitation on total reproductive structures (estimated reproduction) for 14 focal species. Regression slopes are from models that include effects of temperature and collection day-of-year. Five species had significantly less reproduction with more precipitation (red lines), while two species had increased reproduction (blue lines). Plots for temperature and climatic water deficit (CWD) are in Fig. S10. (b and c) Summarized effects of (b) precipitation and temperature (in the same model) and (c) CWD on estimated reproduction. Point-ranges show the mean and 95% credible interval of the slope of the regression of total reproduction on each climate variable. Grey point ranges indicate that the effects of climate are not different from zero. None of these responses to climate showed significant phylogenetic signal (Table S3).

### Q2: Does later onset of seasonal rainfall alter phenology or reduce estimated reproduction?

For nine species, earlier rainfall onset resulted in specimens that were collected at a more advanced phenological stage (with a greater proportion of flowers or fruits) relative to specimens that experienced later rainfall onset (Fig. 2). These effects were present when also taking into account collection timing, average temperature, and total precipitation during the specimens’ growing seasons (Fig. S9). In contrast, with these same covariates in the model, the timing of germination-triggering rains had no effects on estimated specimen reproduction except for one species, *S. drepanoides*, which produced relatively more fruit with later rainfall onset (Figs. 4, S11).

**Figure 4.**
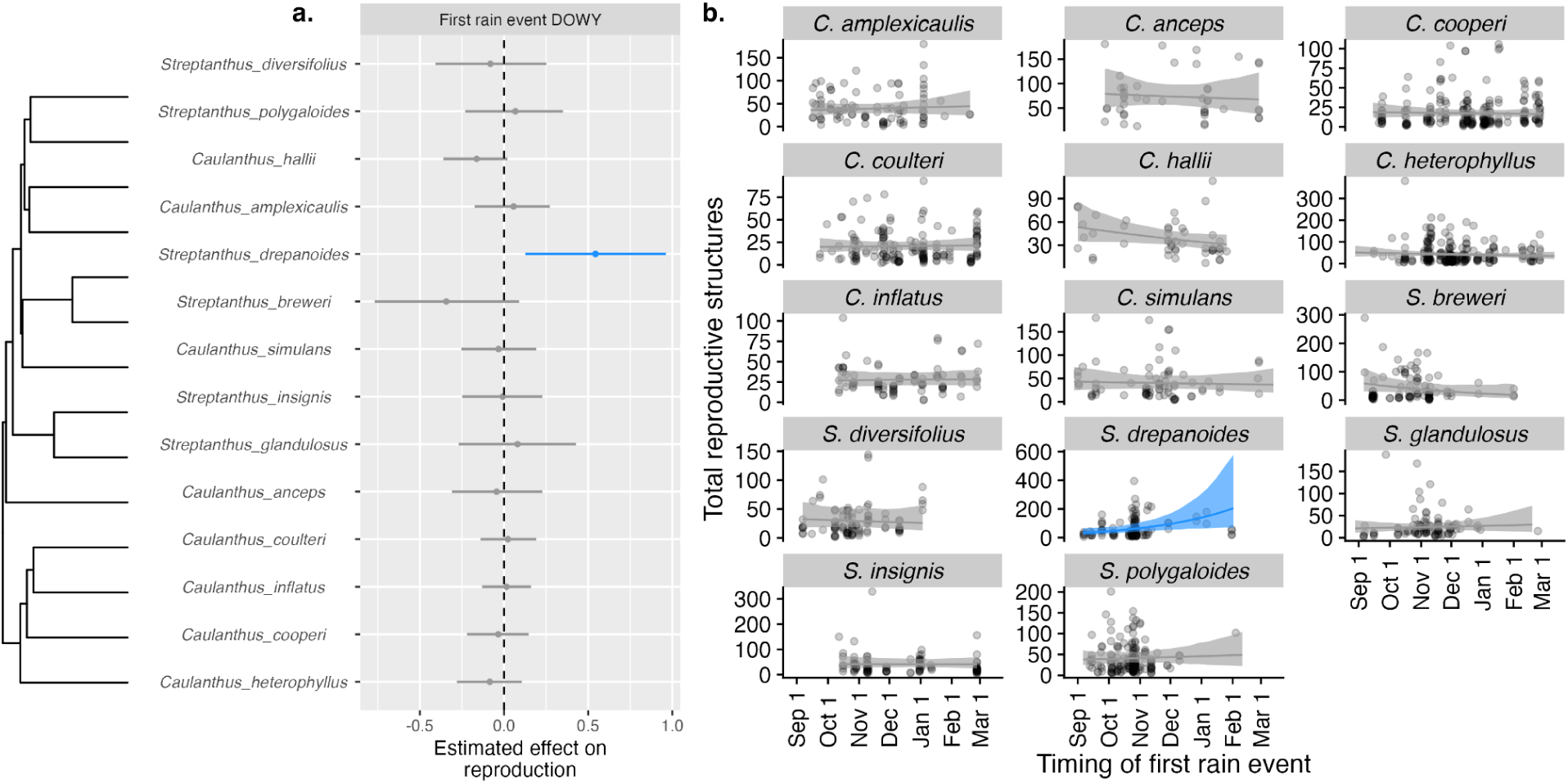
Effects of timing of germination triggering rainfall based on daily precipitation data on estimated reproduction of herbarium specimens. Blue symbols/lines show a positive effect of later germination on reproduction and grey lines are not different from zero. Collection date, total precipitation and average temperature were included in the model as predictors (see Fig. S11). In (b) these slopes are plotted over the raw data points. There was significant phylogenetic signal in the slopes of reproduction responses to the timing of germination-triggering rainfall (first rain event DOWY; Table S3).

### Q3: Across species, are phenological and estimated reproduction responses to climate variables correlated?

Across species, we asked whether species with greater phenological plasticity (i.e., larger effects of climate on phenological stage) had lower fitness sensitivity to climate (i.e., smaller effects of climate on estimated reproduction). Slopes of phenological responses were not significantly related to slopes of total reproduction responses for any climate variable (Fig. S12, Table S2).

### Q4: Are responses to climate evolutionarily conserved across the clade?

We found no evidence of evolutionary history (phylogenetic signal) influencing phenological or estimated reproductive responses to temperature, precipitation, or CWD (Table S3). However, there was significant phylogenetic signal in both phenological and reproductive responses to the timing of first rain events, which are likely to trigger germination (K>1, P<0.001 for phenology, and <0.026 for total reproduction; Figs. 2, 4; Table S3). Phylogenetic signal arises when traits (e.g. slope of relationship between specimen phenology and rainfall timing) of closely related species are more similar to each other than to those of more distantly related species; for example, the subclade of southern *Caulanthus* species share very similar shallow slopes in their phenological responses to rainfall onset timing, and these differ from the magnitude of slopes of the subclade of closely related *S. breweri* and *S. drepanoides,* which share similar responses to rainfall onset (Fig. 2).

## Discussion

Herbarium collections offer incredible resources for understanding the impacts of climate change on plant performance as they span many climatological conditions, large geographic regions and integrate over biotic as well as abiotic environments. A growing number of studies have shown that the phenological stages of herbarium specimens at collection often reflect the climate in which they grew (Primack *et al*., 2004; Miller-Rushing *et al*., 2006; Calinger *et al*., 2013; Jones & Daehler, 2018; DeLeo *et al*., 2020; Love & Mazer, 2021; Pearson *et al*., 2021; Willems *et al*., 2022; Dangremond *et al*., 2022; Ramirez-Parada *et al*., 2024). Our study contributes to this literature by relating climate variables not only to specimen phenology but also to estimated specimen reproduction, which, in annual species, is a key fitness component. Importantly, we also consider climate-performance relationships in light of specimen-specific tailored estimates of climate (see also Bontrager & Angert, 2016; Hereford *et al*., 2017; Love & Mazer, 2021; Mazer *et al*., 2021 for other studies exploring more tailored climate windows). The lived climate of a specimen was characterized from estimated germination date to species average collection date using fine-scaled temporal and spatial climate records and previously gained knowledge of germination cues (Worthy *et al*., 2024). In the course of analyzing data with these aims, we came to the realization that collection protocols aimed at accurate species identification could be supplemented with protocols that would improve our ability to measure climate impacts on plants and population viability using specimen data. Thus, we also propose collection protocol additions that could complement the current practices for specimen collection.

### Effects of lived climate on plant phenology and reproduction

Of four potentially important climate variables (average temperature, total precipitation, timing of precipitation onset, and total CWD), timing of onset and total amount of precipitation had the greatest impacts on plant phenology and reproduction, based on the number of significant relationships and the magnitude of slopes of these relationships (Figs. 1-4). Ten species exhibited a significant relationship between total precipitation and phenological stage, and eight species demonstrated a relationship between total precipitation and total reproduction. In contrast, seven species had relationships with phenology and temperature, and only two showed relationships between temperature and reproduction (Figs. 1, 3). A recent meta-analysis of climate impacts on phenology in field experiments finds temperature to be more important than precipitation in predicting phenological shifts (Zhou et al., 2023); similarly, a meta-analysis of 21 plant traits measured in the field also found temperature to be more important than precipitation in influencing trait values (Moles *et al*., 2014). However, in water-limited ecosystems, precipitation may be the dominating force influencing phenology (see meta-analysis of Lu *et al*., 2023), especially for annual species with less access to deep water reserves (Currier & Sala, 2022). For example, phenological change in grasses in an arid ecosystem was more impacted by precipitation than temperature (Currier & Sala, 2022).

### Lack of correspondence between phenological and total reproduction responses

If climate-sensitive phenology is adaptive, then we expect species that are able to adjust their phenology in response to climate variation to be buffered against effects of climate variation on fitness. In this case, we would expect an overall pattern where large climate effects on phenology are associated with small climate effects on reproduction. Phenological shifts can compensate to some degree for climatic stress, reducing the impact of climate on total reproduction (Martínez-Berdeja *et al*., 2023; Miller & Stuble, 2024; Worthy *et al*., 2025 and see below). In California mediterranean ecosystems, fall precipitation onset, which is the cue for germination in many annual species (e.g., Keeley, 1991; Levine *et al*., 2008; Worthy *et al*., 2024) is occurring later, and providing less water (Swain *et al*., 2018; Luković *et al*., 2021). We found that later rainfall onset resulted in specimens with more delayed phenology in 9 of our 14 study species, after the effects of growing season temperature and precipitation amounts were accounted for. These effects on phenology, however, did not translate into significant effects on estimated reproduction (Figs. 3, 4, S11), suggesting the possibility of compensatory buffering of phenological responses to stabilize fitness responses (Martínez-Berdeja *et al*., 2023; Miller & Stuble, 2024; Worthy *et al*., 2025). Our previous experimental work showed that many of these species can speed phenological phases, for example by accelerating flowering rate in response to later germination timing (Worthy *et al*., 2025), as has also been observed in other annual species in Mediterranean climates (Steyn *et al*., 1996; Olliff-Yang & Ackerly, 2021; Martínez-Berdeja *et al*., 2023).

Whether traits related to climate adaptation are constrained in their evolution may indicate how species adapt to changing climates. In this case, the responses of species to rainfall onset timing in both phenology and total reproduction were the only relationships that were evolutionarily conserved (phenology: Fig. 2, Table S3; reproduction: Fig. 4; Table S3). Moreover, the general pattern of responses, with the southern *Caulanthus* subclade being less responsive phenologically to rainfall (shallower slopes), is consistent with results of our previous watering experiment, in which these species matured quickly and did not extend their flowering season when provided with ample water (Pearse *et al*., 2020). Combined, our results suggest that adaptation to changes in rainfall timing might be more constrained than adaptation to total amounts of rainfall or rising temperatures (see also Bradshaw & Holzapfel, 2006).

Another potential mechanism underlying the lack of correspondence between the effects of climate on phenology and on total reproduction entails interactions with other species (Farías *et al*., 2021; Van Dyke *et al*., 2022; Armitage, 2024; Terry, 2024). These include climate-induced changes in competitive hierarchies (e.g., Niu & Wan, 2008), mismatch or facilitation of pollination (e.g., Kudo & Ida, 2013; Gezon *et al*., 2016; Tiusanen *et al*., 2020; Freimuth *et al*., 2022), changes in interactions with herbivores (Fabina *et al*., 2010; Meineke *et al*., 2018; Hamann *et al*., 2021), or other interactors like microbes (Ware *et al*., 2021). For example, many species in this clade experience strong negative effects of competition from other species (Cacho & Strauss, 2014), which may explain why reproduction is negatively associated with precipitation for several species in our analyses, as competition from other individuals or species is likely higher in wetter years.

On top of all these ecological complexities is the possibility that specimens may not have been collected at an advanced enough phenological stage to provide a good estimate of the true total reproductive potential of the specimen. The average specimen was 28% fruit at time of collection. Despite this limitation, we did find significant effects of precipitation on total reproduction in five species. All of these complexities could reduce the linkages between climate and estimated reproduction in herbarium specimens, even when climate may be impacting plant reproduction.

Additional types of herbarium protocols could make linkages between specimen reproduction and climate more useful for projecting resilience to climate change (see Box 1. *Making collections even more valuable for climate change research*). If we want to use specimens to answer questions like the ones posed here—specifically, assessing fitness responses to ongoing climate change—we suggest new protocols that could make collections even more valuable for these purposes (Box 1). The focus is on adding different types of specimen collections, and not replacing existing ones. Most of our suggestions include targeted increased sampling effort within species and across years. We also highlight possibilities to link to community science digital image databases like iNaturalist. Conclusions Herbarium specimens are valuable resources that preserve a record of the effects of historical conditions on plant traits and performance. Our study contributes to methodological approaches by characterizing climate tailored to the lifespan of the specimen. We achieved this by estimating germination timing of each specimen, based on prior experimental results on germination cues. Our results highlight how herbarium specimens can lend more insight into effects of climate variation on plant performance and revealed that timing of precipitation onset, in particular, is an important climate factor influencing herbarium specimen phenology across this clade. While phenological responses to climate are important, linking those changes to plant fitness will ultimately determine the usefulness of herbarium specimens to inform climate impacts on the viability of plant populations (see also Iler *et al*., 2019). Both phenology and reproduction were generally more affected by precipitation levels than temperature in this water-limited mediterranean system. Our results also suggest that specimen total reproduction, as we have estimated it, appears substantially less affected by climate than is plant phenology, a result which could stem from phenology buffering plant fitness, interactions with other species, or other mechanisms. Ultimately, this buffering may signal robustness of these populations in the face of changing precipitation and temperature regimes, but it may also indicate a lack of good estimates of total plant reproduction from specimens. To increase confidence in estimating climate impacts on plant fitness from herbarium specimens, we suggest additional collection practices (Box 1) and linkages with citizen science databases like iNaturalist (Heberling & Isaac, 2018; Iler *et al*., 2019) focused on place-centered collections sampled intensively through time.

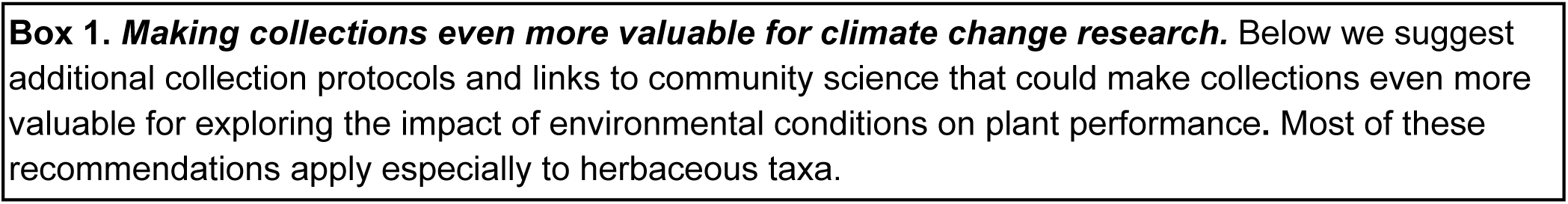
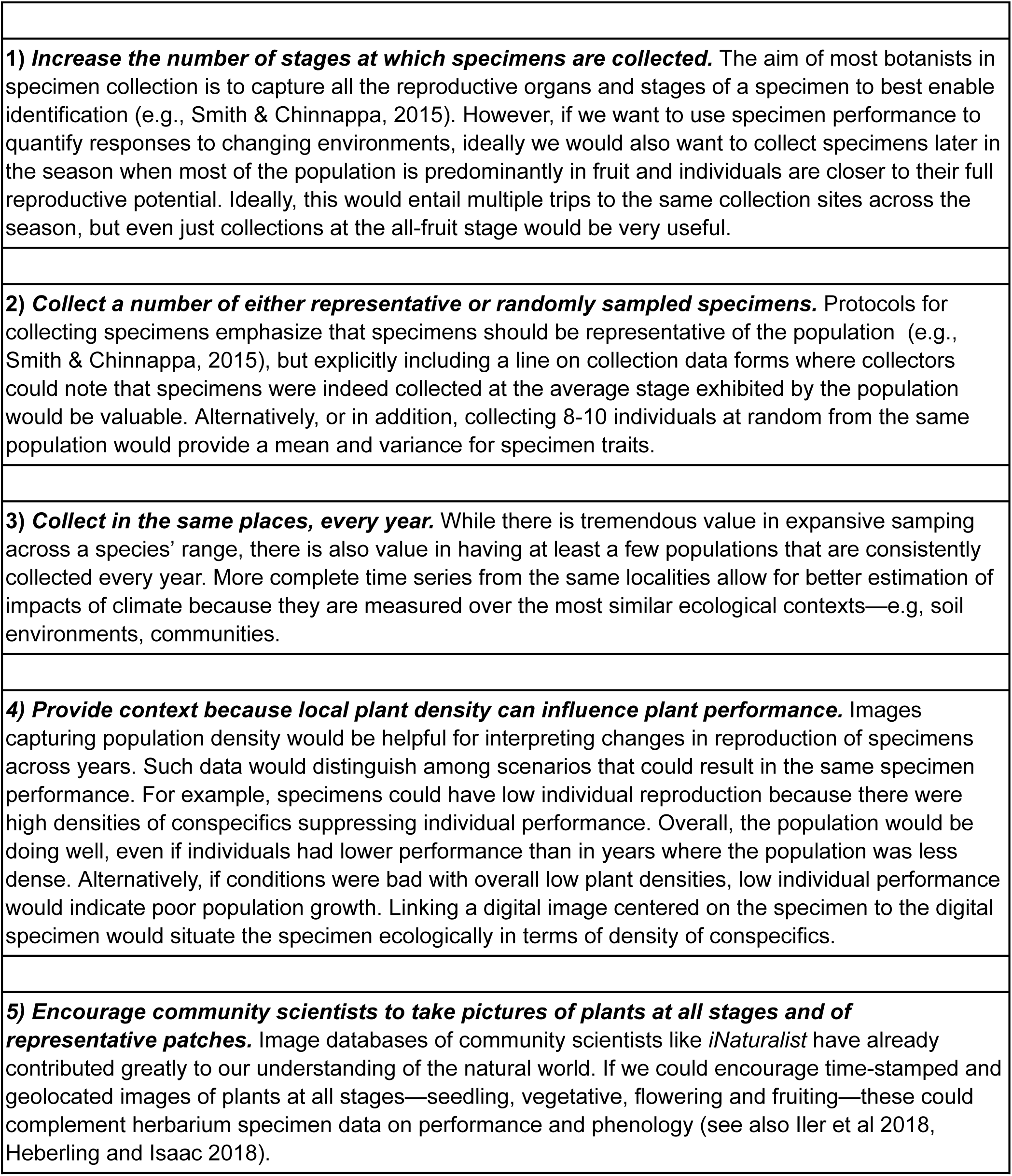

## Acknowledgements

We thank undergraduate assistants Eda Ceviker, Ian Clark, Macarena Cortina Petrasic, Kees Hood, Lara Hsia, Evan Jordan, Louisa Liu, Maya Martinez, Adrianna Ng, Natascha Paxton, Shannon Reilly, Lila Simpson and research technician Elizabeth Davis for data collection on digitized herbarium specimens. We thank the California Academy of Science

Herbarium and the UC Davis Herbarium and acknowledge the amazing data resource that the Consortium of California Herbaria provides (ucjeps.berkeley.edu/consortium/). This work was funded by US National Science Foundation DEB 1831913 to Gremer (PI), Schmitt, Maloof and Strauss.

## Competing interests

No authors have competing interests.

## Author contributions

SYS, MGB, JS, JRG designed the work. MGB and SYS developed species and specimen sampling protocols, MGB and SJW conducted data analyses in substantial consultation with all authors; MGB and LCL oversaw specimen reproductive data collection, JAM contributed to image analysis protocols; SYS and MGB wrote the first draft, with contributions from SJW. All authors commented substantively on the draft.

## Data availability

Upon acceptance all data and code will be publicly archived. All data and code will be made available to reviewers upon request.

---

**Figure S1.**
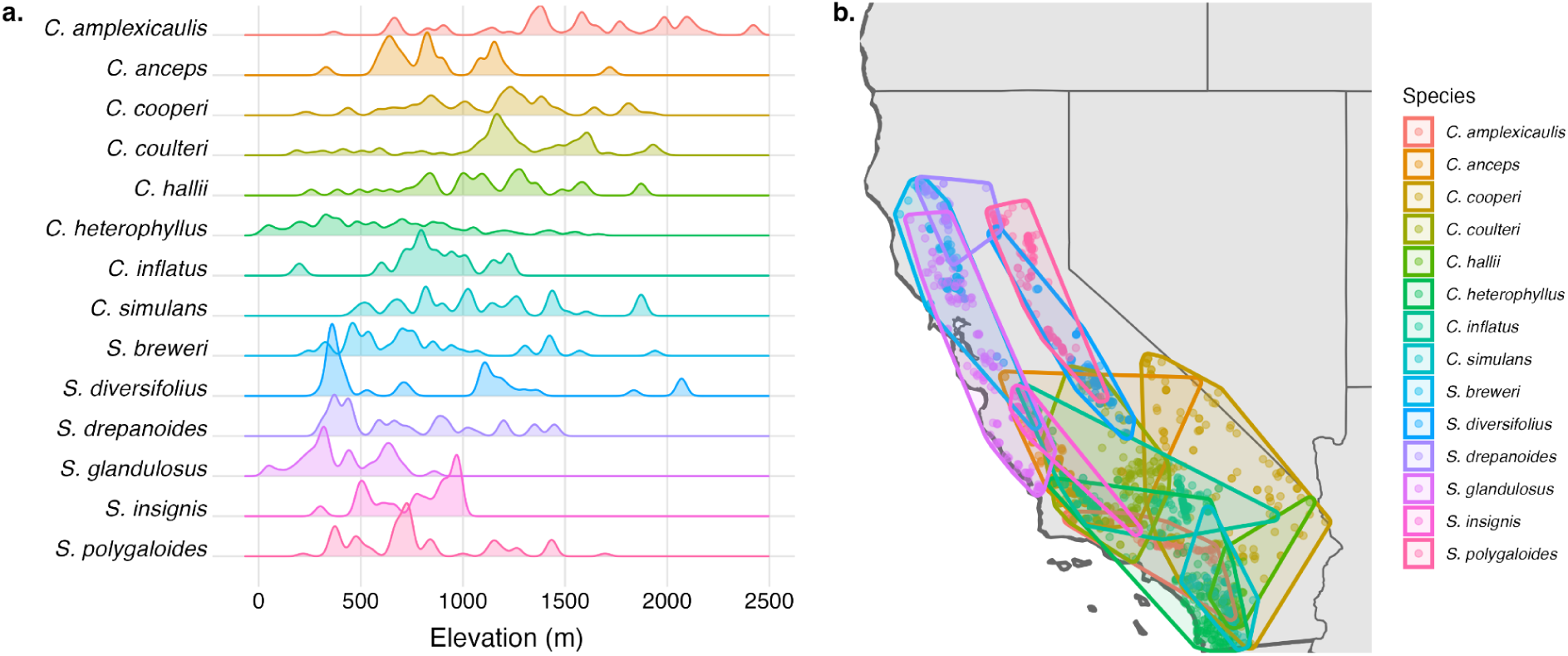
Elevational and geographic distributions for the 14 *Streptanthus* (s.l.) species in our study. (a) Density ridges of elevational data for the 868 specimens in our dataset with elevations recorded; heights of ridges indicate the relative frequencies of specimens from that elevation (relative within species). (b) Locations (points) of the specimens in our dataset, colored by species. Convex hulls are drawn around the specimen locations for each species.

**Figure S2.**
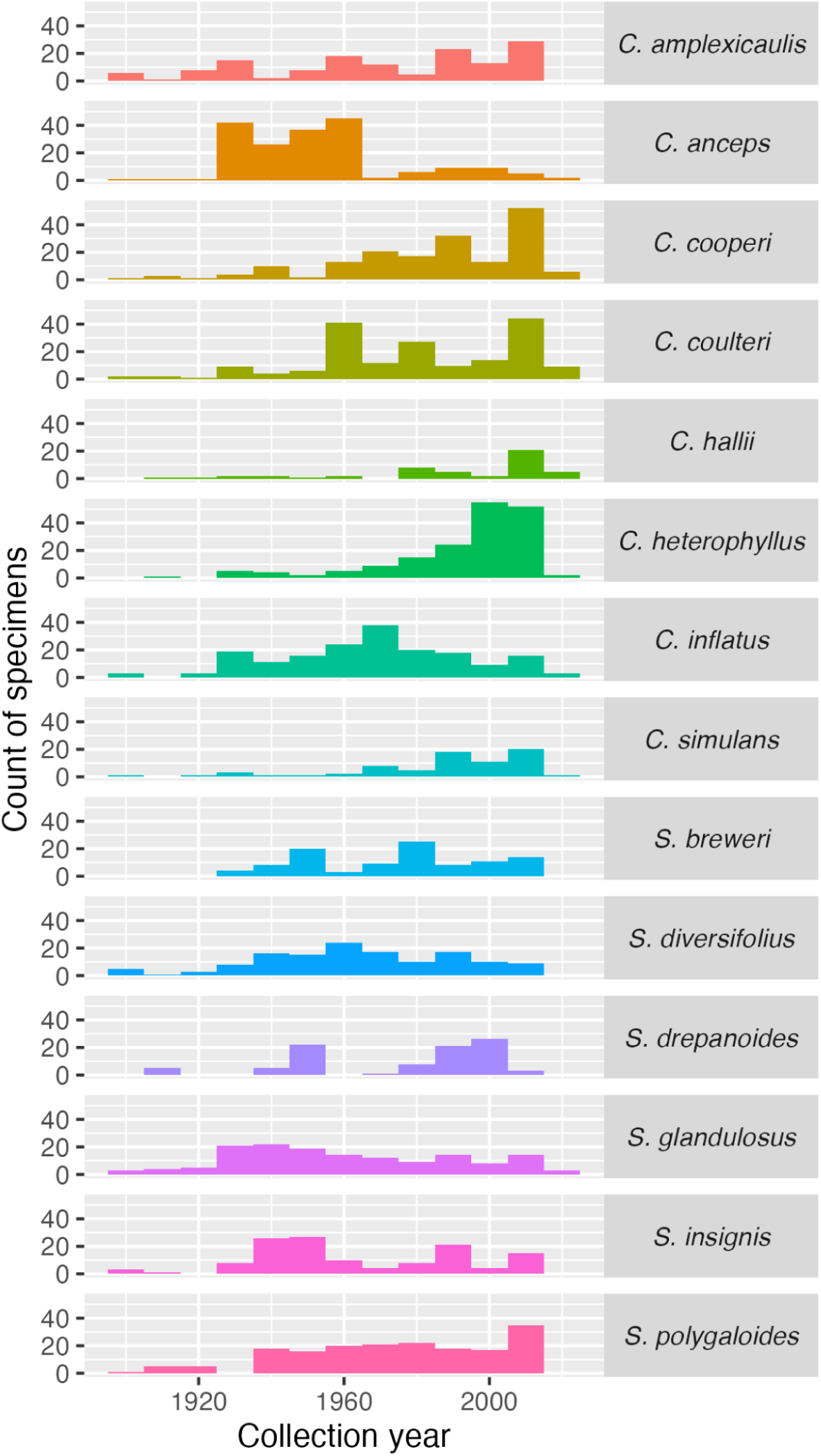
Species included in this study were represented by specimens collected across a range of years. Each bin represents 10 years, beginning with the years 1898-1907.

**Figure S3.**
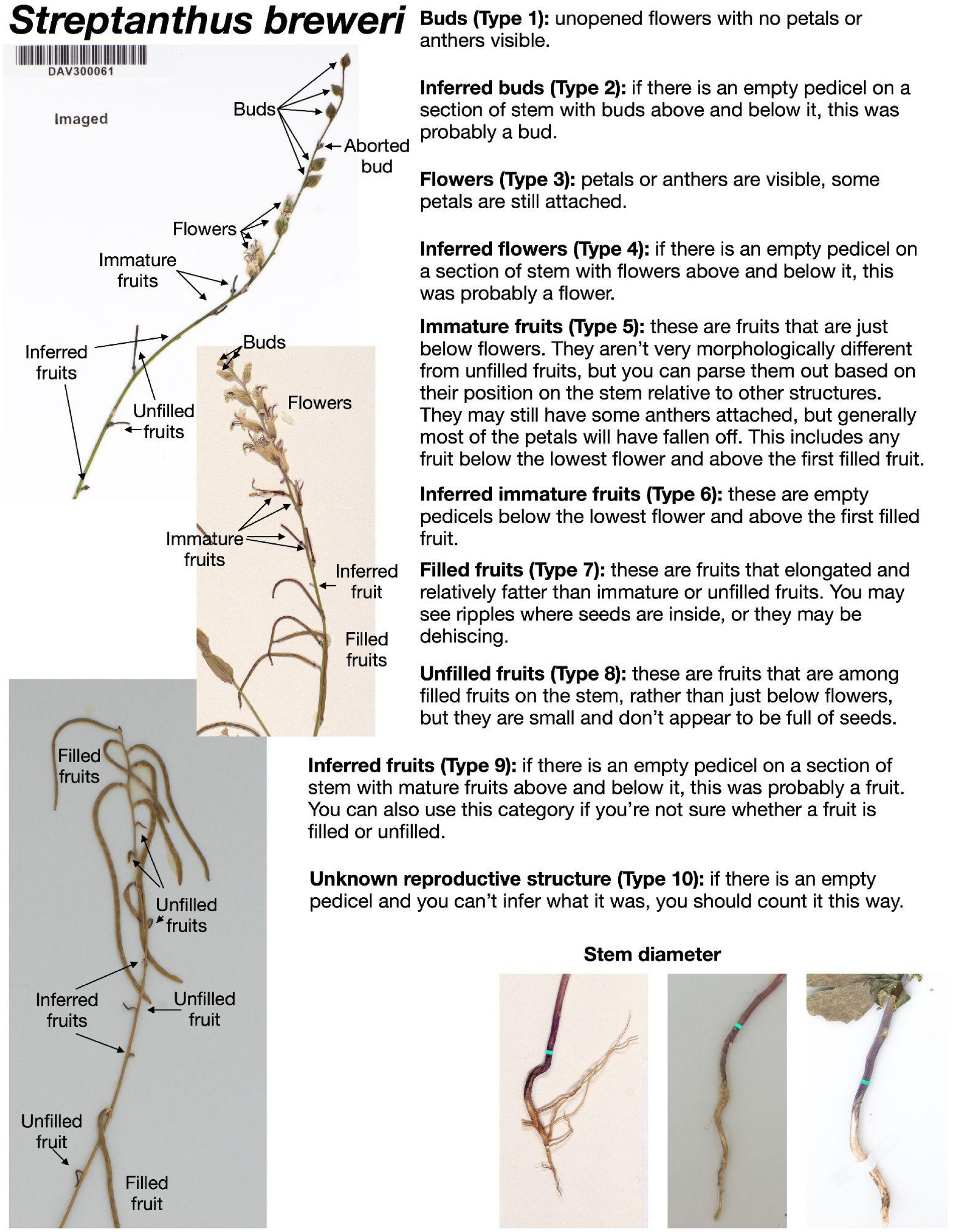
Example guide for counting reproductive structures on herbarium samples; *Streptanthus breweri*.

**Figure S4.**
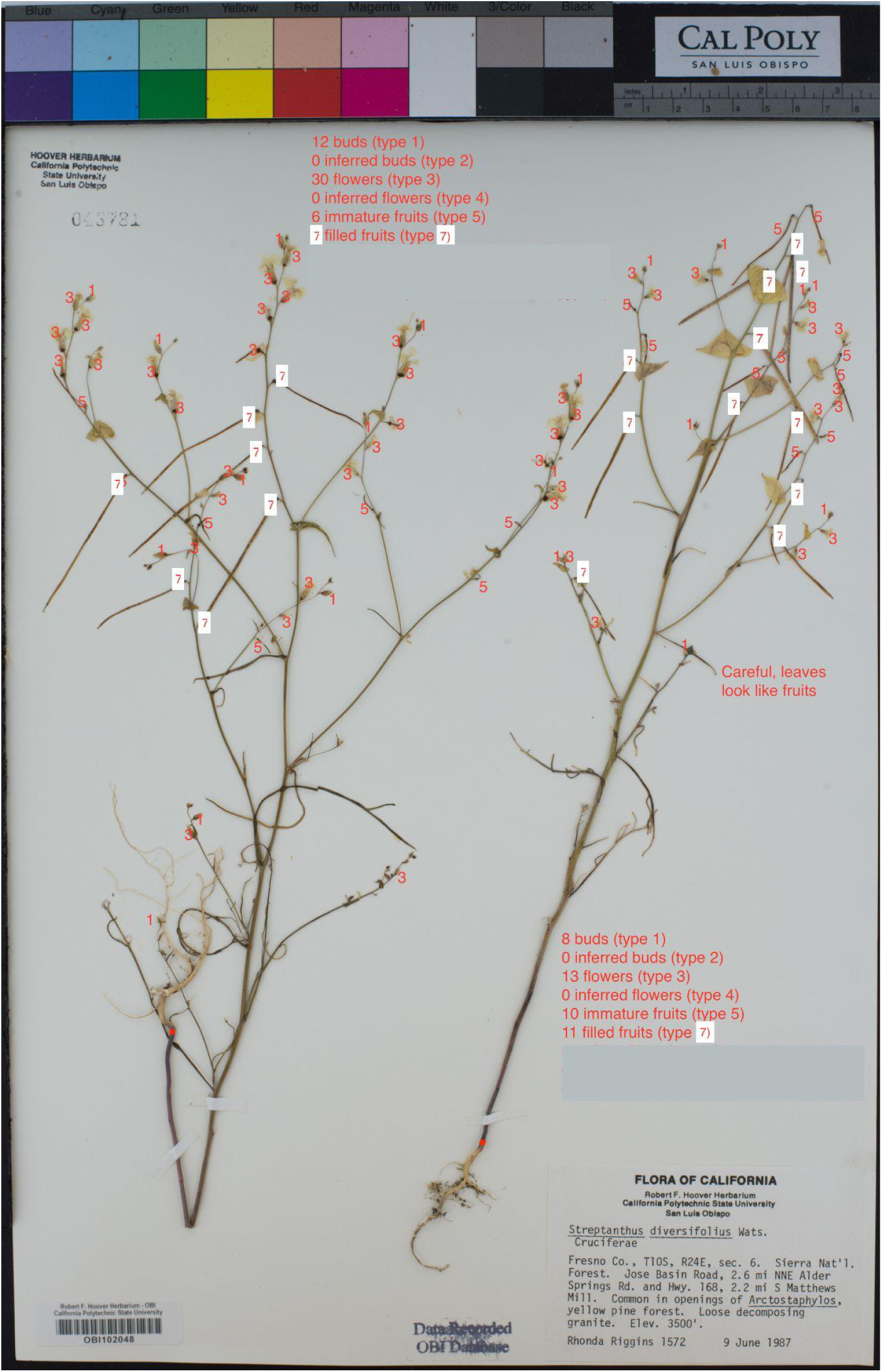
Example of a counted specimen sheet for *S. diversifolius*. Reproductive structures were classified into categories (see text and Fig. S3). Specimens were inspected to make sure they were whole and undamaged by browsers. Note that for this species, leaves on flowering stems can look like fruits and are distinguished from fruits by lacking pedicels.

**Figure S5.**
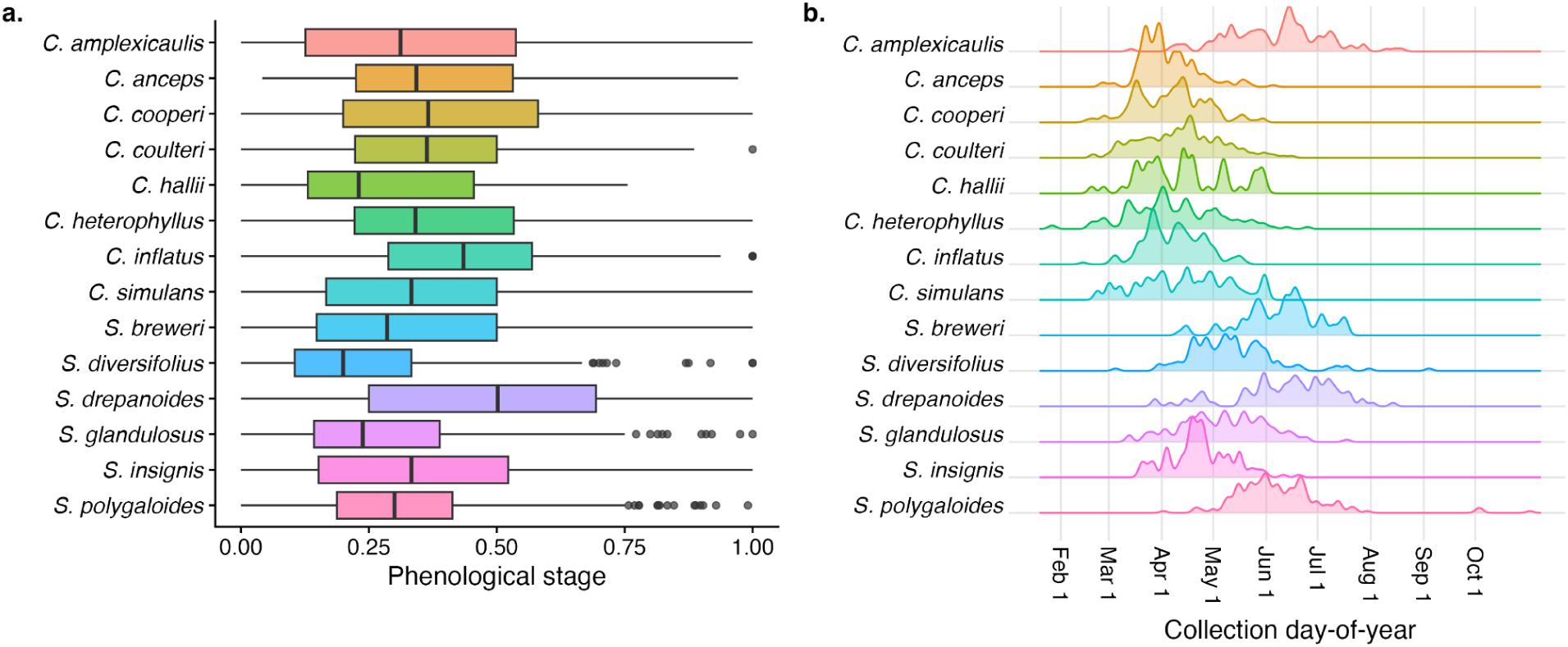
(a) Phenological stages of specimens: these are calculated based on a weighted sum of flowers, immature, and mature fruits, divided by a total count of reproductive structures (including buds). Specimens at a phenological stage of 0 are entirely in bud, those with a phenological stage of 1 have only mature fruits, and intermediate values indicate a combination of buds, flowers, immature fruits, and mature fruits. (b) Specimen collection timing varied within and between species, with the majority of specimens collected in March-June. This panel shows density ridges; heights of ridges indicate the relative frequencies of specimens from that day of year (relative within species).

**Figure S6.**
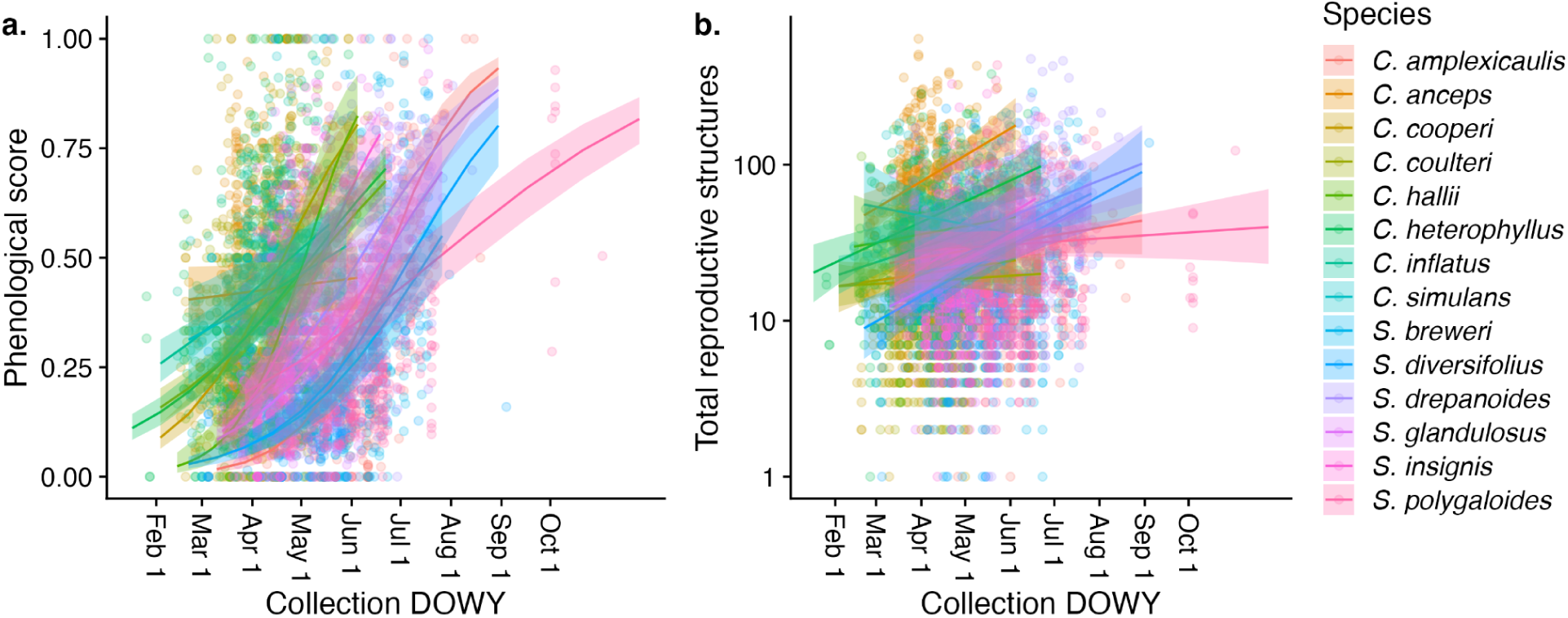
Relationship between collection day-of-year (DOY) and a) phenology and b) estimated reproduction. Collection day-of-year was represented in models as the number of days since September 1 in the year prior to collection until the collection date. In a), Specimens entirely in bud would have a phenological score of 0, and specimens with only mature fruits would have a phenological score of 1. Overall, collection day-of-year was strongly related to phenology, and less so to total reproduction; in each case, there remains much variation that may be explained by climate variables.

**Figure S7.**
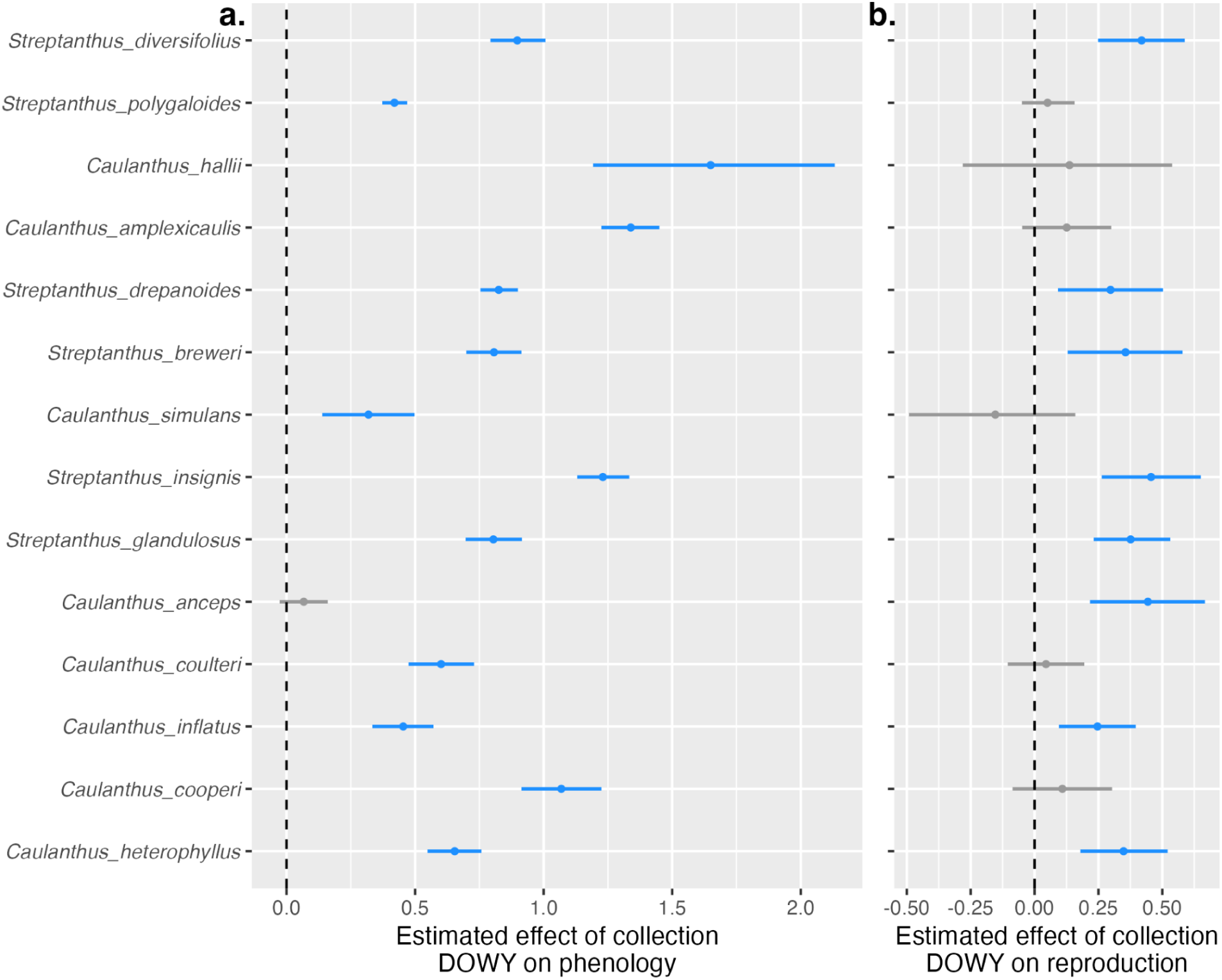
Effects of collection day-of-year covariate on phenology and reproduction. These estimates are from models that also included covariates of temperature and precipitation (Figs. 1, 3). Blue point ranges indicate species for which specimens collected later in the year had significantly (a) more advanced phenology or (b) greater total reproduction relative to specimens collected earlier in the year. Grey point ranges indicate species for which the effect of collection day-of-year on phenology or reproduction was not different from 0; there were no cases where specimens with later collection day-of-year had more delayed phenology or less reproduction. These effects are also shown plotted over the raw data in Fig. S6.

**Figure S8.**
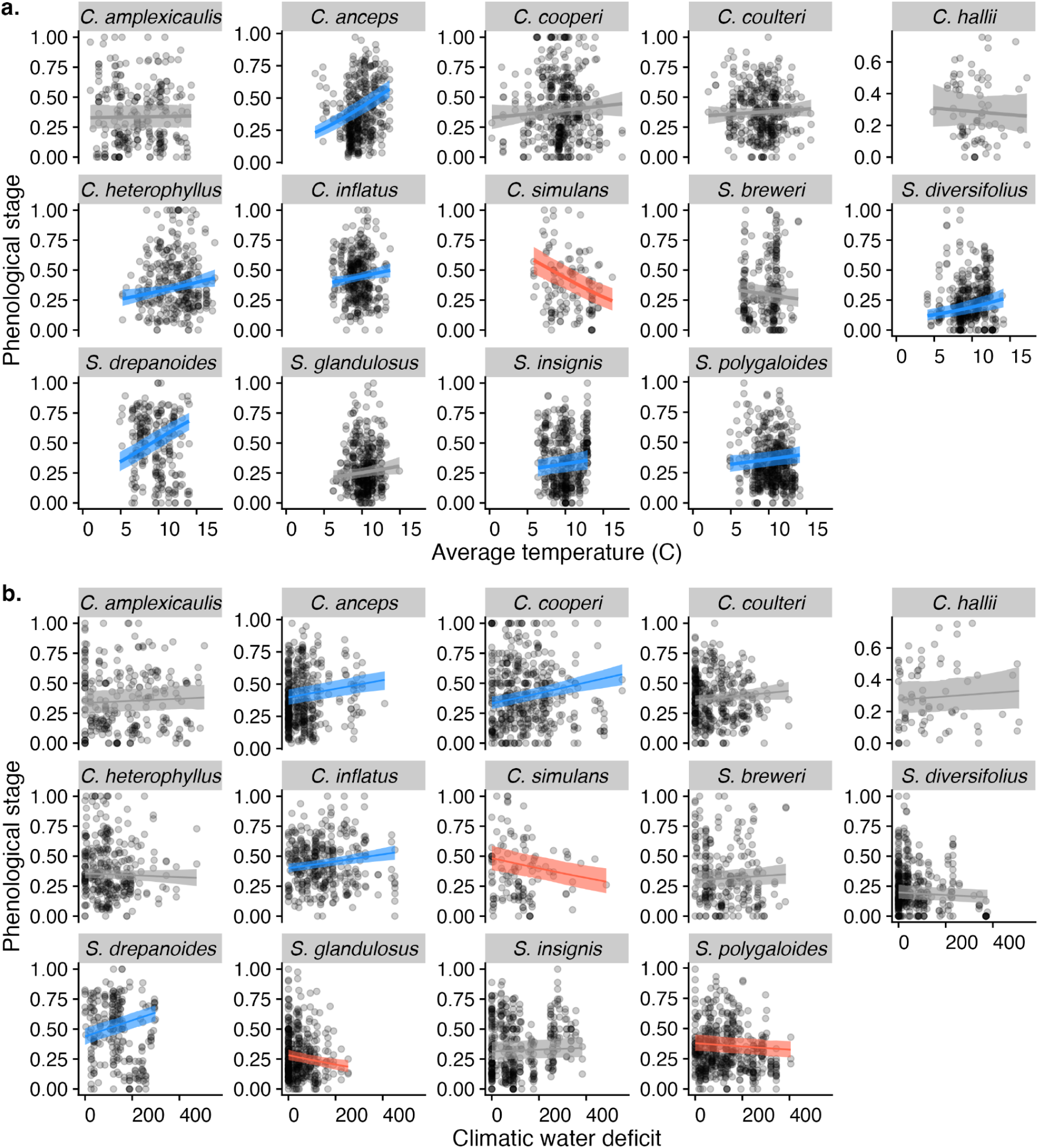
(a) Effects of average temperature on phenological stage for 14 focal species from the *Strepthanthus* clade. One species had significantly less advanced phenology under warm temperatures (red line), while seven species had more advanced phenology under warm temperatures (blue lines). These regression slopes are from models that also include effects of precipitation and collection day-of-year. (b) Effects of total climatic water deficit (CWD) on phenological stage. Four species had significantly more advanced phenology with high CWD (blue lines), while three species had less advanced phenology under high CWD (red lines).Regression slopes are from models that also include effects of collection day-of-year.

**Figure S9.**
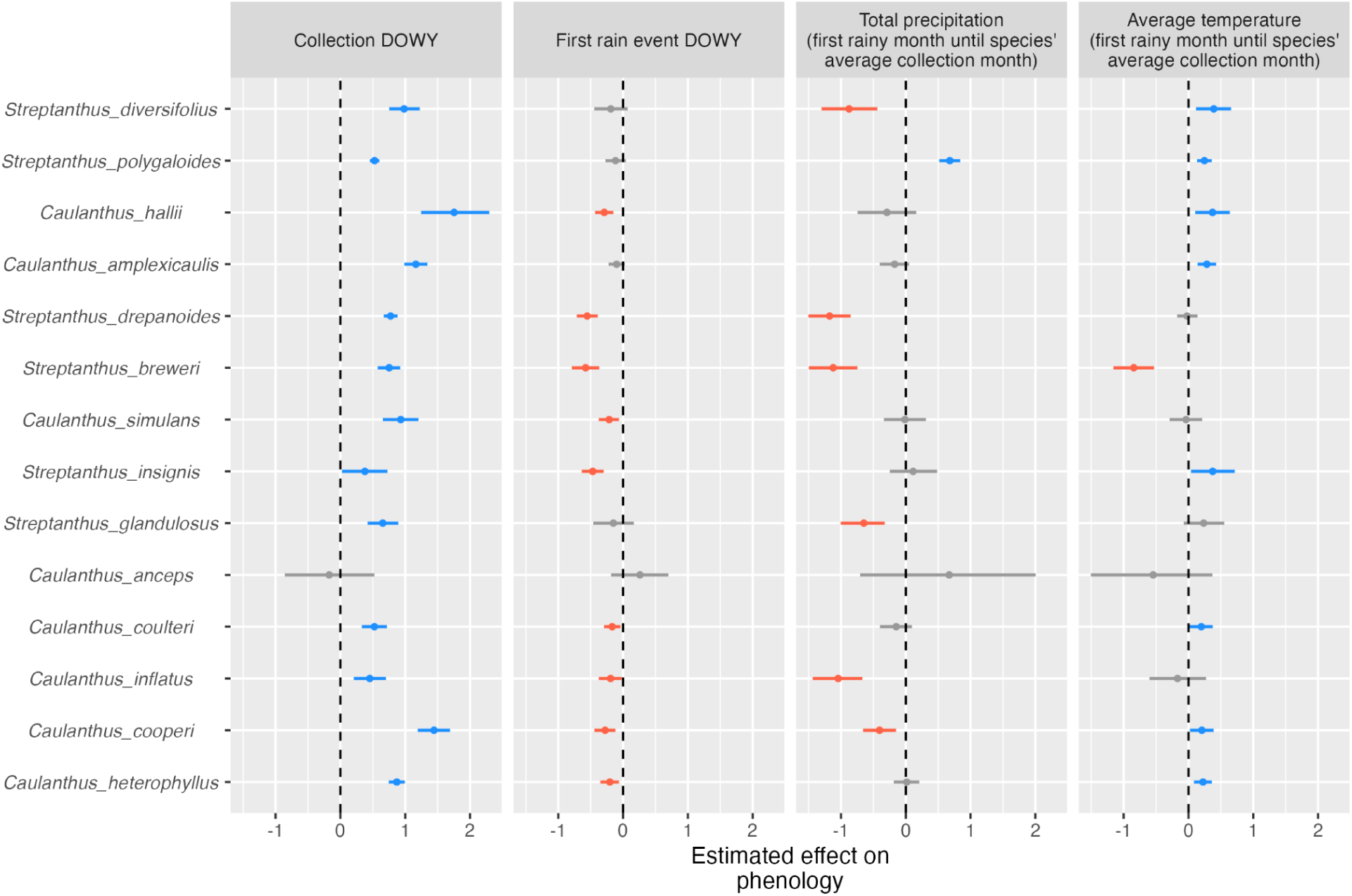
Summarized effects of collection day-of-year, timing of first rain event, precipitation, and temperature on specimen phenological scores. Point-ranges show the mean and 95% credible interval of the slope of the regression of phenological score on each variable. Blue point-ranges indicate that higher values of the variable advanced phenology, red point-ranges indicate that higher values of the variable delayed phenology, and grey point ranges indicate where effects are not different from zero. These results are from a model using only specimens for which daily precipitation data was available—a smaller dataset than we used in our primary analysis of temperature and precipitation effects.

**Figure S10.**
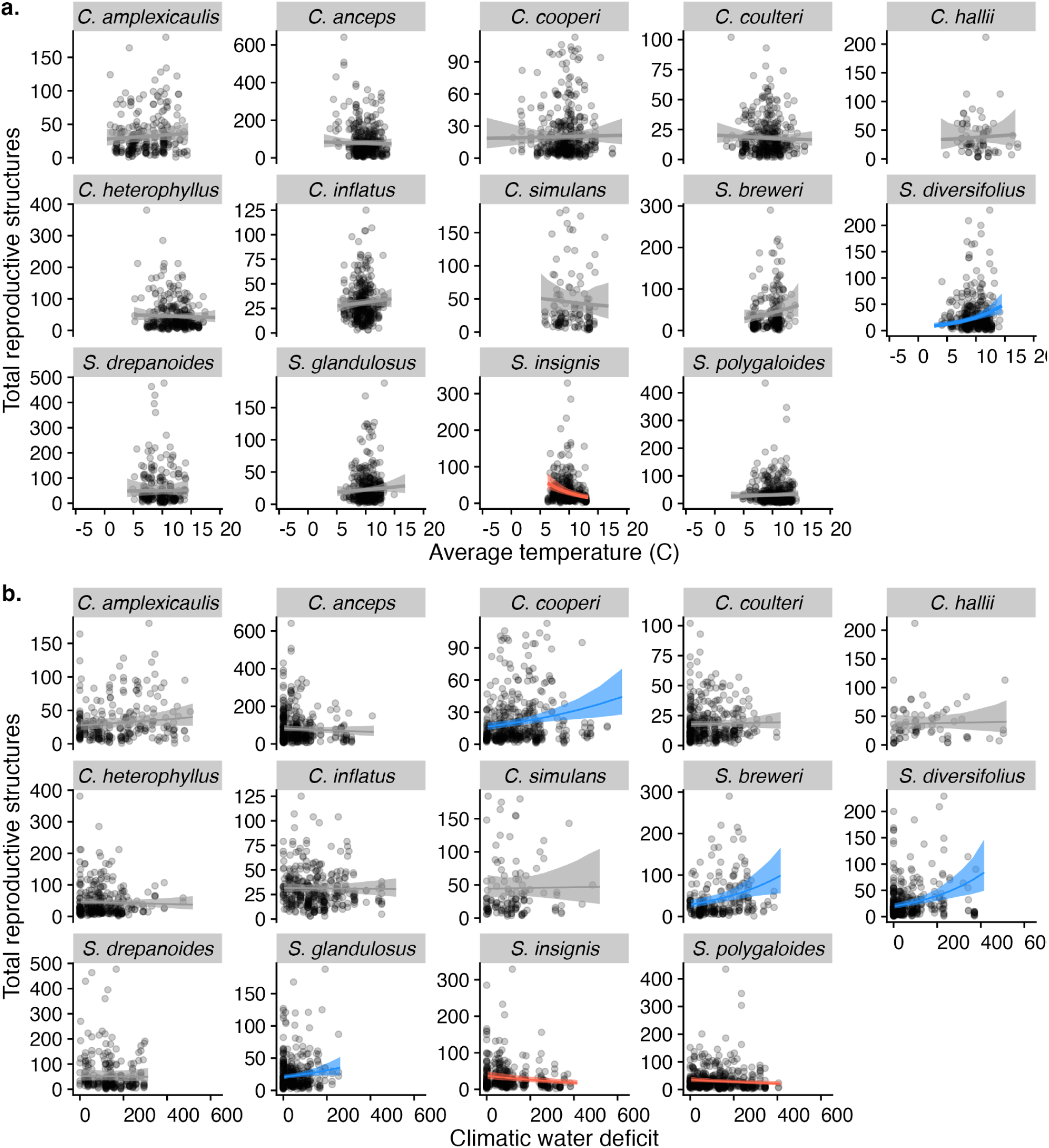
(a) Effects of average temperature on total reproductive structures for 14 focal species from the *Strepthanthus* clade. Regression slopes are from models that also include effects of precipitation and collection day-of-year. One species had significantly less reproduction withunder warm temperatures (red line), while one species had increased reproduction under warm temperatures (blue line). (b) Effects of climatic water deficit (CWD) on total reproductive structures. Regression slopes are from models that also include effects of collection day-of-year. Four species had significantly more reproduction with high CWD (blue lines), while two species had decreased reproduction with high CWD (red lines).

**Figure S11.**
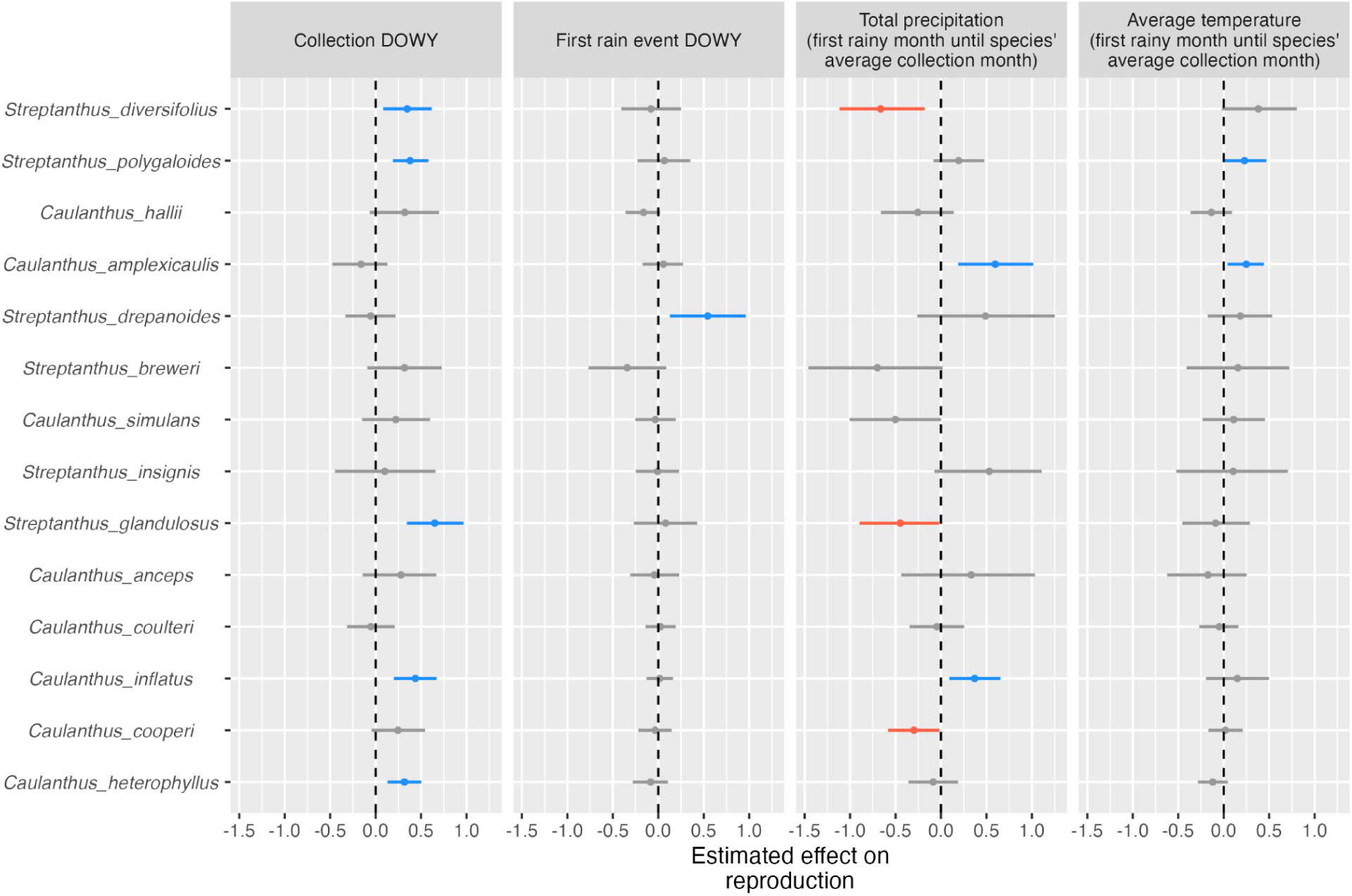
Summarized effects of collection day-of-year, timing of first rain event, precipitation, and temperature on the number of reproductive structures on each specimen. Point-ranges show the mean and 95% credible interval of the slope of the regression of reproduction on each variable. Blue point-ranges indicate that higher values of the variable are associated with increased reproduction, red point-ranges indicate that higher values of the variable are associated with decreased reproduction, and grey point ranges indicate where effects are not different from zero. These results are from a model using only specimens for which daily precipitation data was available—a smaller dataset than we used in our primary analysis of temperature and precipitation effects.

**Figure S12.**
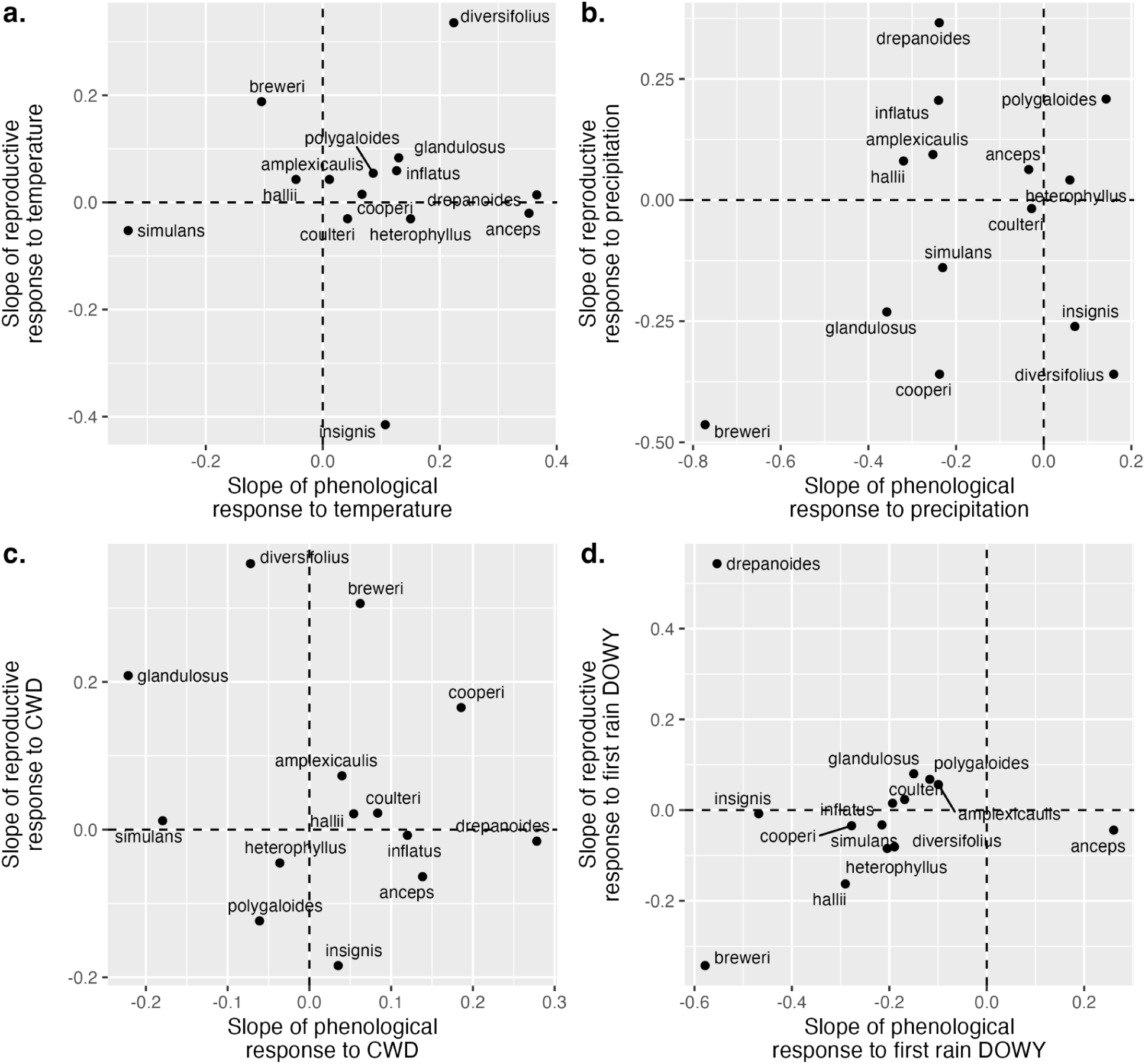
Scatterplots relating how species responded phenologically to climate versus how they responded in total reproduction to the same to climate variables; these relationships are asking whether there is any systematic across-species correlation in how species respond phenologically and reproductively: (a) temperature, (b) total precipitation, (c) total climatic water deficit (CWD), and (d) the day-of-year of the first rainfall event. Slopes in (a) are also shown in Figs. 1b, S8a, 3b and S10a. Slopes in (b) are also shown in Figs. 1ab and 3ab. Slopes in (c) are also shown in Figs. 1c, S8b, 1c and S10b. Slopes in (d) are also shown in Figs. 2 and 4. We found no significant association between phenological and reproductive responses for these climate variables (Table S2).

**Table S1.** Categorization rules of reproductive structures on herbarium specimens.

| Category | Description |
| --- | --- |
| Buds | unopened flowers with no petals visible |
| Inferred buds | contextual placement indicates a bud (i.e., buds are above and below), but only the pedicel remains |
| Flowers | anthers are visible, petals are still attached |
| Inferred flowers | contextual placement indicates a flower, but only the pedicel remains |
| Immature fruits | styles which have not yet widened and elongated with seeds; these occur in the section of the stem below flowers |
| Inferred immature fruits | contextual placement indicates an immature fruit, but only the pedicel remains |
| Filled fruit | fruit has elongated and swelling from seeds may be visible |
| Unfilled fruits | fruits that have not elongated or swollen with seeds; morphologically similar to immature fruits but distinguished by their context (filled fruits higher on the stem indicate that these have had time to mature) |
| Inferred fruits | contextual placement indicates either a filled or unfilled fruit, but only the pedicel remains |
| Unknown reproductive structure | a pedicel is present, but no structure is present, and there is not adequate context to assume what type of structure it was |

**Table S2.** Results from phylogenetically independent contrasts between phenological and reproductive effort response slopes for four climate variables.

| Variable | Estimate | SE | t-value | P-value |
| --- | --- | --- | --- | --- |
| Temperature | 0.091 | 0.29 | 0.31 | 0.76 |
| Precipitation | 0.22 | 0.34 | 0.66 | 0.52 |
| Climatic water deficit | -0.19 | 0.31 | -0.61 | 0.55 |
| First rain event DOWY | -0.021 | 0.25 | -0.085 | 0.93 |

**Table S3.** Tests for phylogenetic signal in the effect sizes of regressions of estimated reproduction (“reproduction”) and phenology (“phenological stage”) to climate and collection date. We report Blomberg’s K and estimate the significance of phylogenetic signal using 10,000 randomizations where effect sizes were randomized across the tips of the phylogeny and K was repeatedly calculated—this significance is reported in the P-value column. When P < 0.05, we further test whether the estimate of K was significantly >1 by comparing our estimate of K to a null distribution of 10,000 K values generated under Brownian motion, a null model of evolution. Values of K significantly >1 indicate that species tend to have more similar phenology and reproduction responses to climate than expected based on a Brownian motion model of evolution, i.e. more evolutionary constraint. DOWY = day of water year (days counted from September 1). All models also included a random effect of collection location (see methods).

| Variable | K | P-value | Notes |
| --- | --- | --- | --- |
| *Model: Reproduction ~ average temperature + log(total precipitation) + collection DOWY* |  |  |  |
| Average temperature | 0.895 | 1 |  |
| log(Total precipitation) | 0.886 | 0.481 |  |
| *Model: Phenological stage ~ average temperature + log(total precipitation) + collection DOWY* |  |  |  |
| Average temperature | 0.797 | 0.846 |  |
| log(Total precipitation) | 0.860 | 0.745 |  |
| *Model: Reproduction ~ total climatic water deficit (CWD) + collection DOWY* |  |  |  |
| Total CWD | 0.812 | 0.915 |  |
| *Model: Phenological stage ~ total climatic water deficit (CWD) + collection DOWY* |  |  |  |
| Total CWD | 0.961 | 0.266 |  |
| *Model: Reproduction ~ first rain event DOWY + average temperature + log(total precipitation) + collection DOWY* |  |  |  |
| **First rain event DOWY** | **1.77** | **0.0001** | **Tested if significantly $>1$ by comparison to null distribution. P-value $< 0.001$, so significantly greater than 1.** |
| Average temperature | 0.765 | 0.234 |  |
| log(Total precipitation) | 0.891 | 0.573 |  |
| *Model: Phenological stage ~ first rain event DOWY + average temperature + log(total precipitation) + collection DOWY* |  |  |  |
| **First rain event DOWY** | **1.310** | **0.028** | **Tested if significantly $>1$ by comparison to null distribution. P-value = 0.022, so significantly greater than 1.** |
| Average temperature | 1.049 | 0.452 |  |
| log(Total precipitation) | 1.004 | 0.323 |  |

## Notes

### Competing Interest Statement

The authors have declared no competing interest.

### Summary of Updates

We have revised the abstract, clarified our explanation of the phenological index, and updated the section of the discussion on herbarium collection methods, in addition to small clarifications throughout the manuscript.

## References

1. Armitage DW. 2024. To remain modern the coexistence program requires modern statistical rigour. Nature 632: E15–E20.

2. Barnett KL, Facey SL. 2016. Grasslands, Invertebrates, and Precipitation: A Review of the Effects of Climate Change. Frontiers in Plant Science 7.

3. Beatley JC. 1974. PHENOLOGICAL EVENTS AND THEIR ENVIRONMENTAL TRIGGERS IN MOJAVE-DESERT ECOSYSTEMS. Ecology 55: 856–863.

4. Blomberg SP, Garland T, Ives AR. 2003. TESTING FOR PHYLOGENETIC SIGNAL IN COMPARATIVE DATA: BEHAVIORAL TRAITS ARE MORE LABILE. Evolution 57: 717–745.

5. Bontrager M, Angert AL. 2016. Effects of range-wide variation in climate and isolation on floral traits and reproductive output of Clarkia pulchella. American Journal of Botany 103: 10–21.

6. Bontrager M, Worthy SJ, Cacho NI, Leventhal L, Maloof JN, Gremer JR, Schmitt J, Strauss SY. 2025. Herbarium specimens reveal a constrained seasonal climate niche despite diverged annual climates across a wildflower clade. : 2025.02.28.640808.

7. Bradshaw WE, Holzapfel CM. 2006. Evolutionary Response to Rapid Climate Change. Science 312: 1477–1478.

8. Bürkner P-C. 2017. brms: An R Package for Bayesian Multilevel Models Using Stan. Journal of Statistical Software 80: 1–28.

9. Cacho NI, McIntyre PJ, Kliebenstein DJ, Strauss SY. 2021. Genome size evolution is associated with climate seasonality and glucosinolates, but not life history, soil nutrients or range size, across a clade of mustards. Annals of Botany 127: 887–902.

10. Cacho NI, Millie Burrell A, Pepper AE, Strauss SY. 2014. Novel nuclear markers inform the systematics and the evolution of serpentine use in Streptanthus and allies (Thelypodieae, Brassicaceae). Molecular Phylogenetics and Evolution 72: 71–81.

11. Cacho NI, Strauss SY. 2014. Occupation of bare habitats, an evolutionary precursor to soil specialization in plants. Proceedings of the National Academy of Sciences 111: 15132–15137.

12. Calinger KM, Queenborough S, Curtis PS. 2013. Herbarium specimens reveal the footprint of climate change on flowering trends across north-central North America. Ecology Letters 16: 1037–1044.

13. Currier CM, Sala OE. 2022. Precipitation versus temperature as phenology controls in drylands. Ecology 103: e3793.

14. Dangremond EM, Hill CH, Louaibi S, Muñoz I. 2022. Phenological responsiveness and fecundity decline near the southern range limit of Trientalis borealis (Primulaceae). Plant Ecology 223: 41–52.

15. Davis CC, Willis CG, Connolly B, Kelly C, Ellison AM. 2015. Herbarium records are reliable sources of phenological change driven by climate and provide novel insights into species’ phenological cueing mechanisms. American Journal of Botany 102: 1599–1609.

16. DeLeo VL, Menge DNL, Hanks EM, Juenger TE, Lasky JR. 2020. Effects of two centuries of global environmental variation on phenology and physiology of *Arabidopsis thaliana*. Global Change Biology 26: 523–538.

17. Di Castri F, Mooney HA. 2012. Mediterranean Type Ecosystems: Origin and Structure. Springer Science & Business Media.

18. Fabina NS, Abbott KC, Gilman RT. 2010. Sensitivity of plant–pollinator–herbivore communities to changes in phenology. Ecological Modelling 221: 453–458.

19. Farías AA, Armas C, Gaxiola A, Cea AP, Luis Cortés J, López RP, Casanoves F, Holmgren M, Meserve PL, Gutiérrez JR, et al.2021. Species interactions across trophic levels mediate rainfall effects on dryland vegetation dynamics. Ecological Monographs 91: e01441.

20. Flint LE, Flint AL. 2014. California Basin Characterization Model: A dataset of historical and future hydrologic response to climate change. USGS data release Version 1.1.

21. Flint L, Flint A, Thorne J, Boynton R. 2013. Fine-scale hydrologic modeling for regional landscape applications: the California Basin Characterization Model development and performance. Ecological Processes 2: 25.

22. Freimuth J, Bossdorf O, Scheepens JF, Willems FM. 2022. Climate warming changes synchrony of plants and pollinators. Proceedings of the Royal Society B: Biological Sciences 289: 20212142.

23. Gezon ZJ, Inouye DW, Irwin RE. 2016. Phenological change in a spring ephemeral: implications for pollination and plant reproduction. Global Change Biology 22: 1779–1793.

24. González-Megías A, Menéndez R. 2012. Climate change effects on above- and below-ground interactions in a dryland ecosystem. Philosophical Transactions of the Royal Society B: Biological Sciences 367: 3115–3124.

25. Gremer JR, Chiono A, Suglia E, Bontrager M, Okafor L, Schmitt J. 2020. Variation in the seasonal germination niche across an elevational gradient: the role of germination cueing in current and future climates. American Journal of Botany 107: 350–363.

26. Hamann E, Wadgymar SM, Anderson JT. 2021. Costs of reproduction under experimental climate change across elevations in the perennial forb Boechera stricta. Proceedings of the Royal Society B: Biological Sciences 288: 20203134.

27. Hänel S, Tielbörger K. 2015. Phenotypic response of plants to simulated climate change in a long-term rain-manipulation experiment: a multi-species study. Oecologia 177: 1015–1024.

28. Hatchett BJ, Koshkin AL, Guirguis K, Rittger K, Nolin AW, Heggli A, Rhoades AM, East AE, Siirila-Woodburn ER, Brandt WT, et al.2023. Midwinter Dry Spells Amplify Post-Fire Snowpack Decline. Geophysical Research Letters 50: e2022GL101235.

29. Heberling JM, Isaac BL. 2018. iNaturalist as a tool to expand the research value of museum specimens. Applications in Plant Sciences 6: e01193.

30. Hereford J, Schmitt J, Ackerly DD. 2017. The seasonal climate niche predicts phenology and distribution of an ephemeral annual plant, Mollugo verticillata. Journal of Ecology 105: 1323–1334.

31. Iler AM, Compagnoni A, Inouye DW, Williams JL, CaraDonna PJ, Anderson A, Miller TEX. 2019. Reproductive losses due to climate change-induced earlier flowering are not the primary threat to plant population viability in a perennial herb. Journal of Ecology 107: 1931–1943.

32. Jones CA, Daehler CC. 2018. Herbarium specimens can reveal impacts of climate change on plant phenology; a review of methods and applications. PeerJ 6: e4576.

33. Keeley JE. 1991. Seed germination and life history syndromes in the California chaparral. The Botanical Review 57: 81–116.

34. Kudo G, Ida TY. 2013. Early onset of spring increases the phenological mismatch between plants and pollinators. Ecology 94: 2311–2320.

35. Levine JM, McEachern AK, Cowan C. 2008. Rainfall effects on rare annual plants. Journal Of Ecology 96: 795–806.

36. Levine JM, McEachern AK, Cowan C. 2011. Seasonal timing of first rain storms affects rare plant population dynamics. Ecology 92: 2236–2247.

37. Love NLR, Bonnet P, Goëau H, Joly A, Mazer SJ. 2021. Machine Learning Undercounts Reproductive Organs on Herbarium Specimens but Accurately Derives Their Quantitative Phenological Status: A Case Study of Streptanthus tortuosus. Plants 10: 2471.

38. Love NLR, Mazer SJ. 2021. Region-specific phenological sensitivities and rates of climate warming generate divergent temporal shifts in flowering date across a species’ range. American Journal of Botany 108: 1873–1888.

39. Love NLR, Park IW, Mazer SJ. 2019. A new phenological metric for use in pheno-climatic models: A case study using herbarium specimens of *Streptanthus tortuosus*. Applications in Plant Sciences 7: e11276.

40. Lu C, Zhang J, Min X, Chen J, Huang Y, Zhao H, Yan T, Liu X, Wang H, Liu H. 2023. Contrasting responses of early- and late-season plant phenophases to altered precipitation. Oikos 2023: e09829.

41. Lüdecke D. 2018. ggeffects: Tidy Data Frames of Marginal Effects from Regression Models. Journal of Open Source Software 3: 772.

42. Luković J, Chiang JCH, Blagojević D, Sekulić A. 2021. A Later Onset of the Rainy Season in California. Geophysical Research Letters 48: e2020GL090350.

43. Martínez-Berdeja A, Okada M, Cooper MD, Runcie DE, Burghardt LT, Schmitt J. 2023. Precipitation timing and soil substrate drive phenology and fitness of Arabidopsis thaliana in a Mediterranean environment. Functional Ecology 37: 2471–2487.

44. Mazer SJ, Love NLR, Park IW, Ramirez-Parada T, Matthews ER. 2021. PHENOLOGICAL SENSITIVITIES TO CLIMATE ARE SIMILAR IN TWO CLARKIA CONGENERS: INDIRECT EVIDENCE FOR FACILITATION, CONVERGENCE, NICHE CONSERVATISM, OR GENETIC CONSTRAINTS. Madroño 68.

45. Meineke EK, Davis CC, Davies TJ. 2018. The unrealized potential of herbaria for global change biology. Ecological Monographs 88: 505–525.

46. Miller CN, Stuble KL. 2024. Warm Spring Days are Related to Shorter Durations of Reproductive Phenophases for Understory Forest Herbs. Ecology and Evolution 14: e70700.

47. Miller-Rushing AJ, Primack RB, Primack D, Mukunda S. 2006. Photographs and herbarium specimens as tools to document phenological changes in response to global warming. American Journal of Botany 93: 1667–1674.

48. Moles AT, Perkins SE, Laffan SW, Flores-Moreno H, Awasthy M, Tindall ML, Sack L, Pitman A, Kattge J, Aarssen LW, et al.2014. Which is a better predictor of plant traits: temperature or precipitation? Journal of Vegetation Science 25: 1167–1180.

49. Niu S, Wan S. 2008. Warming changes plant competitive hierarchy in a temperate steppe in northern China. Journal of Plant Ecology 1: 103–110.

50. Olliff-Yang RL, Ackerly DD. 2021. LATE PLANTING SHORTENS THE FLOWERING PERIOD AND REDUCES FECUNDITY IN LASTHENIA CALIFORNICA. Madroño 68: 377–387.

51. Paradis E, Schliep K. 2019. ape 5.0: an environment for modern phylogenetics and evolutionary analyses in R. Bioinformatics 35: 526–528.

52. Pathak TB, Maskey ML, Dahlberg JA, Kearns F, Bali KM, Zaccaria D. 2018. Climate Change Trends and Impacts on California Agriculture: A Detailed Review. Agronomy 8: 25.

53. Pearse IS, Aguilar JM, Strauss SY. 2020. Life-History Plasticity and Water-Use Trade-Offs Associated with Drought Resistance in a Clade of California Jewelflowers. The American Naturalist 195: 691–704.

54. Pearse IS, McIntyre P, Cacho NI, Strauss SY. 2022. Fitness homeostasis across an experimental water gradient predicts species’ geographic range and climatic breadth. Ecology 103: e3827.

55. Pearson KD, Love NLR, Ramirez-Parada T, Mazer SJ, Yost JM. 2021. PHENOLOGICAL TRENDS IN THE CALIFORNIA POPPY (ESCHSCHOLZIA CALIFORNICA): DIGITIZED HERBARIUM SPECIMENS REVEAL INTRASPECIFIC VARIATION IN THE SENSITIVITY OF FLOWERING DATE TO CLIMATE CHANGE. Madroño 68: 343–359.

56. Primack D, Imbres C, Primack RB, Miller-Rushing AJ, Del Tredici P. 2004. Herbarium specimens demonstrate earlier flowering times in response to warming in Boston. American Journal of Botany 91: 1260–1264.

57. Ramirez-Parada TH, Park IW, Record S, Davis CC, Ellison AM, Mazer SJ. 2024. Plasticity and not adaptation is the primary source of temperature-mediated variation in flowering phenology in North America. Nature Ecology & Evolution 8: 467–476.

58. Raven PH, Axelrod DI. 1978. Origin and Relationships of the California Flora. University of California Press.

59. Revell LJ. 2012. phytools: an R package for phylogenetic comparative biology (and other things). Methods in Ecology and Evolution 3: 217–223.

60. Revell LJ, Harmon LJ. 2022. Phylogenetic Comparative Methods in R. Princeton University Press.

61. Schwinning S, Sala OE. 2004. Hierarchy of responses to resource pulses in and and semi-arid ecosystems. Oecologia 141: 211–220.

62. Smith B, Chinnappa CC. 2015. Plant Collection, Identification, and Herbarium Procedures. In: Yeung ECT, Stasolla C, Sumner MJ, Huang BQ, eds. Plant Microtechniques and Protocols. Cham: Springer International Publishing, 541–572.

63. Steyn HM, van Rooyen N, van Rooyen MW, Theron GK. 1996. The phenology of Namaqualand ephemeral species. The effect of water stress. Journal of Arid Environments 33: 49–62.

64. Swain DL, Langenbrunner B, Neelin JD, Hall A. 2018. Increasing precipitation volatility in twenty-first-century California. Nature Climate Change 8: 427–433.

65. Terry JCD. 2024. Uncertain competition coefficients undermine inferences about coexistence. Nature 632: E9–E14.

66. Tevis Jr. L. 1958. Germination and Growth of Ephemerals Induced by Sprinkling a Sandy Desert. Ecology 39: 681–688.

67. Tiusanen M, Kankaanpää T, Schmidt NM, Roslin T. 2020. Heated rivalries: Phenological variation modifies competition for pollinators among arctic plants. Global Change Biology 26: 6313–6325.

68. Van Dyke MN, Levine JM, Kraft NJB. 2022. Small rainfall changes drive substantial changes in plant coexistence. Nature 611: 507–511.

69. Wade RN, Karley AJ, Johnson SN, Hartley SE. 2017. Impact of predicted precipitation scenarios on multitrophic interactions. Functional Ecology 31: 1647–1658.

70. Ware IM, Van Nuland ME, Yang ZK, Schadt CW, Schweitzer JA, Bailey JK. 2021. Climate-driven divergence in plant-microbiome interactions generates range-wide variation in bud break phenology. Communications Biology 4: 1–9.

71. Went FW. 1949. Ecology of Desert Plants. II. The Effect of Rain and Temperature on Germination and Growth. Ecology 30: 1–13.

72. Willems FM, Scheepens JF, Bossdorf O. 2022. Forest wildflowers bloom earlier as Europe warms: lessons from herbaria and spatial modelling. New Phytologist 235: 52–65.

73. Willis CG, Ellwood ER, Primack RB, Davis CC, Pearson KD, Gallinat AS, Yost JM, Nelson G, Mazer SJ, Rossington NL, et al.2017. Old Plants, New Tricks: Phenological Research Using Herbarium Specimens. Trends in Ecology & Evolution 32: 531–546.

74. Worthy SJ, Ashlock SR, Miller A, Maloof JN, Strauss SY, Gremer JR, Schmitt J. 2025. Accelerated Phenology Fails to Buffer Fitness Loss from Delayed Rain Onset in a Clade of Wildflowers. The American Naturalist 205: 485–501.

75. Worthy SJ, Miller A, Ashlock SR, Ceviker E, Maloof JN, Strauss SY, Schmitt J, Gremer JR. 2024. Germination responses to changing rainfall timing reveal potential climate vulnerability in a clade of wildflowers. Ecology n/a: e4423.

76. Yim C, Bellis ES, DeLeo VL, Gamba D, Muscarella R, Lasky JR. 2024. Climate biogeography of *Arabidopsis thaliana* : Linking distribution models and individual variation. Journal of Biogeography 51: 560–574.

77. Zettlemoyer MA, Conner RJ, Seaver MM, Waddle E, DeMarche ML. 2024. A Long-Lived Alpine Perennial Advances Flowering under Warmer Conditions but Not Enough to Maintain Reproductive Success. The American Naturalist: E000–E000.

78. Zhou H, Min X, Chen J, Lu C, Huang Y, Zhang Z, Liu H. 2023. Climate warming interacts with other global change drivers to influence plant phenology: A meta-analysis of experimental studies. Ecology Letters 26: 1370–1381.

